# VHL loss enables immune checkpoint blockade therapy by boosting type I interferon response

**DOI:** 10.1101/2023.11.28.569047

**Authors:** Meng Jiao, Xuhui Bao, Mengjie Hu, Dong Pan, Xinjian Liu, Jonathan Kim, Fang Li, Chuan-Yuan Li

**Affiliations:** Department of Dermatology, Duke University Medical Center, Durham, NC 27710; Department of Biochemistry, Molecular Cancer Research Center, School of Medicine, Sun Yat-sen University, Shenzhen, Guangdong, China; School of Medicine, Duke University, Durham, NC 27710; Department of Pharmacology and Cancer Biology, Duke University Medical Center, Durham, NC 27710; Duke Cancer Institute, Duke University Medical Center, Durham, NC 27710

**Author notes:** Correspondence: Chuan-Yuan Li, PhD Duke University Medical Center Durham, NC 27710.

## Abstract

Despite a moderate mutation burden, clear cell renal cell carcinoma (ccRCC) responds well to immune checkpoint blockade (ICB) therapy. Here we report that loss-of-function mutations in the von Hippel-Lindau (VHL) gene, the most frequent in ccRCC, underlies its responsiveness to ICB therapy. We demonstrate that genetic knockout of the *VHL* gene enhanced the efficacy of anti-PD-1 therapy in multiple murine tumor models in a T cell-dependent manner. Mechanistically, we discovered that upregulation of HIF1α and HIF2α induced by VHL gene loss decreased mitochondrial outer membrane potential and caused the cytoplasmic leakage of mitochondrial DNA (mtDNA), which triggered cGAS-STING activation and induced type I interferons. Our study thus provided novel mechanistic insights into the role of VHL gene loss in potentiating ccRCC immunotherapy.

## Introduction

VHL (von Hippel-Lindau) is an E3 ligase responsible for regulating the cellular levels of the hypoxia-inducible factors (HIFs)^1–3^, which are critical transcriptional factors regulating the cellular response to low oxygen tension^4,5^. Furthermore, as a recognized tumor suppressor, its mutations can cause high HIF activities and increased VEGF levels that promote tumor formation. Indeed, loss of *VHL* is a driver mutation in up to 91% of sporadic ccRCCs, with 55% being loss-of-function (LOF) mutations, including frameshift, nonsense, or small deletions^6–9^. However, the inactivation of VHL alone is not sufficient for RCC development ^10–13^. Mutations in additional genes, such as *PBRM1, BAP1*, and *SETD2*, contribute to tumor development and progression^7,14–17^.

Immune checkpoint blockade (ICB) therapy has recently become a powerful weapon in the expanding arsenal against metastatic ccRCC^6^. ICB therapy has demonstrated superior clinical benefits relative to targeted therapies alone in ccRCC in adjuvant and advanced disease settings ^18–23^. However, we do not have a good mechanistic understanding of why ccRCC responds well to ICB therapy. The current paradigm of whether a particular malignancy responds to cancer immunotherapy primarily focuses on its tumor mutation burden (TMB)^24–26^. The main reasoning is that higher TMB leads to higher numbers of neoantigens likely to be recognized by the host’s T cells, critical for the success of ICB therapy ^27^. This paradigm provides a plausible explanation of why some cancers, such as lung cancer and melanoma, are more responsive to ICB than others. However, it can not explain the responsiveness of ccRCC to ICB therapy: the TMB of ccRCC is only very moderate ^24^. For this reason, there have been many efforts to identify factors responsible for ccRCC responsiveness to ICB therapy ^28–35^. However, no consistent mechanism has emerged to explain the TMB-independent response of ccRCC to ICB therapy. In the present study, we examined if deficiencies in the VHL gene play any role in ccRCC responsiveness to ICB therapy.

## Results

### Vhl deficiency increased susceptibility to ICB therapy in murine tumor models

To evaluate the relevance of *Vhl* gene loss in ICB therapy, we conducted anti-PD1 (αPD1) therapy in several *Vhl*-deficient tumor lines following the schedule depicted in Fig. S1D. In the murine renal cell carcinoma Renca tumor model, αPD1 treatment had minimal effects in control tumors but synergized with *Vhl* gene knockout (**Fig. S1A**), significantly suppressing tumor growth (**Fig. 1A**) and extending the survival of tumor-bearing mice (**Fig. 1B**). Indeed, 2/5 mice in the αPD1-treated *Vhl*-KO group remained long-term survivors, indicating significant synergy between *Vhl* gene loss and ICB therapy. In the B16F10 melanoma line, *Vhl* gene loss (**Fig. S1B**) also significantly attenuated tumor growth in immunocompetent C56BL/6J mice (**Fig. 1C**). Moreover, αPD1 treatment significantly suppressed the growth of *Vhl-KO* tumors (**Fig. 1C-D**), with 1/6 mice remaining tumor-free in the entire period of observation (**Fig. 1D**). We further examined the efficacy of ICB therapy in *Vhl-KO* MC38 tumors (**Fig. S1C**). Our data indicated that αPD1 treatment caused additional growth delay in *Vhl-KO* MC38 tumors, which were already substantially slower than control tumors (**Fig. 1E-F**). Notably, 3/5 mice in the αPD1-treated *Vhl-KO* group were long-term survivors (**Fig. 1F**). To rule out the possibility that the off-target effects of CRISPR/Cas9 were responsible for the behavior of the *Vhl-KO* tumors, we restored *Vhl* expression by ectopic gene transduction (**Fig. S1E**). Our data showed that ectopic expression of wild-type mouse *Vhl (mVhl)* in *Vhl-KO* MC38 cells abrogated the delay in tumor formation from the latter (**Fig. S1F-S1G**), thereby supporting an essential role for VHL in this process. Our results, therefore, suggest that *VHL* loss significantly attenuated tumor growth and made them more susceptible to ICB therapy in multiple murine tumor models.

**Figure 1.**
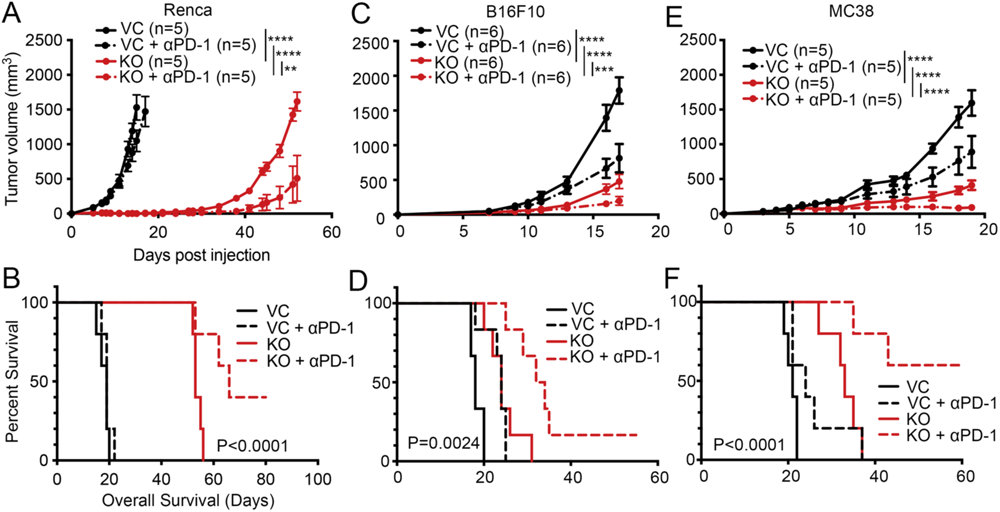
*VHL* loss enhances murine tumor responses to ICB therapy. **(A-B)** Tumor growth (A) and Kaplan-Meier analysis (B) of Balb/c mice subcutaneously inoculated with 1x106 VC or *Vhl*-KO Renca cells (n=5) and treated with isotype control or an αPD-1 antibody. About 100 μg isotype control or αPD-1 antibodies were injected i.p. on at 5, 8, and 11 days post tumor cell inoculation. **(C-D)** Tumor growth (C) and Kaplan-Meier survival analysis (D) of C57BL/6J mice subcutaneously inoculated with 1x105 VC or *Vhl*-KO B16F10 cells (n=5) and treated isotype control or an αPD-1 antibody. About 100 μg isotype control or αPD-1 antibodies were injected i.p. on days 7, 10, 13, and 16 post tumor cell inoculation. **(E-F)** Tumor growth (E) and Kaplan-Meier analysis (F) of C57BL/6J mice subcutaneously inoculated with 5x105 VC or *Vhl*-KO MC38 cells (n=5) and treated with isotype control or αPD-1 antibodies. About 100 μg isotype control or αPD-1 antibodies were injected i.p. on Day 6, 9, and 12 post tumor cell inoculation. Error bars represents mean ± SEM. *P<0.05; **P<0.01; ***P<0.001; ****P<0.0001; n.s. not significant.

### VHL deficiency stimulates intratumoral lymphocyte infiltration

The substantial tumor growth delay caused by *VHL* gene loss and its synergy with αPD1 treatment led us to posit that *VHL* played a critical role in regulating the tumor immune microenvironment. We used flow cytometry to analyze intratumoral infiltration of lymphocytes in control and *Vhl-KO* MC38 tumors to examine our hypothesis (**Fig. S2A**). Our results show significantly more CD3^+^ T cells in *VHL-KO* than in control MC38 tumors (**Fig. 2A**). The increase was observed in both CD4^+^ T-and CD8^+^ T-subsets (**Fig. 2B-C**). Notably, both the numbers of GZMB^+^ CD8^+^ T cells and IFNγ^+^ CD8^+^ T cells were significantly increased (**Fig. 2D-E**), indicating heightened activation of cytotoxic T lymphocytes (CTLs) in *Vhl-KO* tumors. Our analysis also showed high NK^+^ cells in Vhl-KO tumors that did not reach statistical significance (**Fig. 2F**). In contrast, the numbers of F4-80^+^ macrophages (Mý), ψοTCR^+^ T cells, and Foxp3^+^ Tregs were comparable between control and *Vhl-KO* tumors (**Fig. 2G-I**).

**Figure 2.**
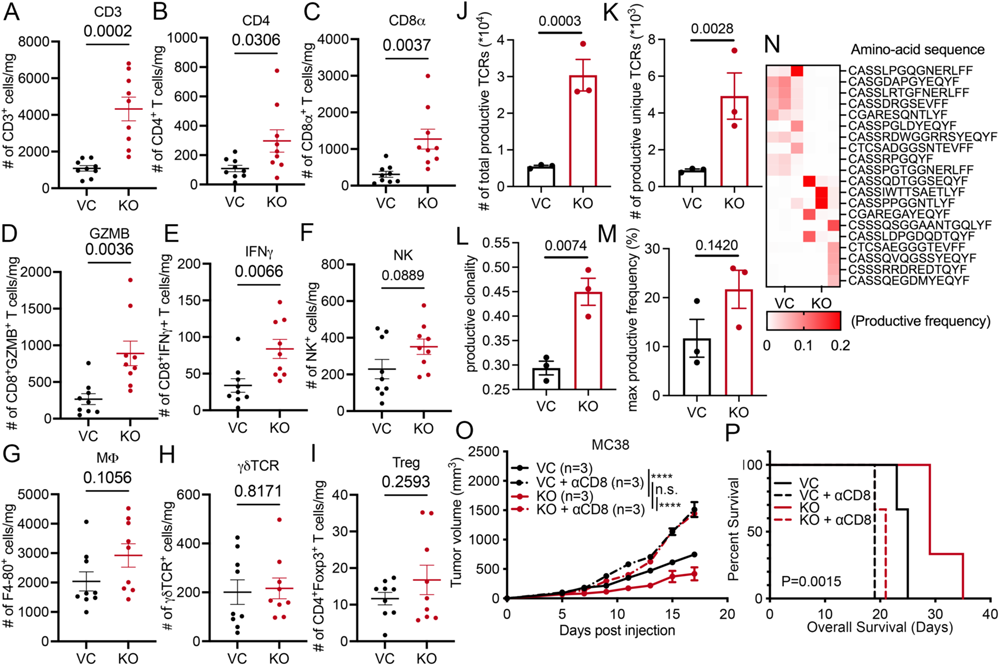
*VHL* loss increases intratumoral lymphocyte infiltration. **(A-I)** Flow cytometry profiling intratumoral lymphocytes in of control and *VHL*-KO MC38 tumors grown in syngeneic C57BL/6 mice. Mice bearing VC or *Vhl*-KO MC38 tumors (n=9) were euthanized on day 15 post tumor cell inoculation. The numbers of immune effector cells per mg of tumor was then obtained using flow cytometry. Error bars represents mean ± SEM. **(J-N)** ImmunoSEQ analysis of TCRβ repertoire. Total productive TCRs (J), productive unique TCRs (K), productive clonality (L), and max productive frequency (M) were shown for control and *Vhl*-KO tumors. Heatmap (N) showing the top 10 (5%) most frequent productive TCR sequences in VC and *Vhl*-KO tumors. **(O-P)** *In vivo* tumor growth and Kaplan-Meier survival analysis of C57BL/6J mice bearing VC and *Vhl*-KO MC38 tumors following depletion of CD8+ T cells using anti-CD8 antibodies. Each mouse received i.p. injection of 100 μg isotype or αCD8b antibodies on Day 1, 4, and 7 post tumor cell inoculation. Error bars represents mean ± SEM. *P<0.05; **P<0.01; ***P<0.001; ****P<0.0001; n.s. not significant.

To further characterize the effect of *Vhl-KO* on intratumoral T cells, we analyzed the T-cell receptor (TCR) repertoire of control and *Vhl*-KO MC38 tumors using the ImmunoSEQ approach ^36^. Our analysis indicated that the numbers of both total productive TCRs (**Fig. 2J**) and productive unique TCRs (**Fig. 2K**) were significantly increased in *Vhl-KO* tumors, suggesting substantially more T cells with increased TCR diversity. Furthermore, productive clonality, an index representing the diversity of rearranged T cell clones, increased substantially in *Vhl-KO* MC38 tumors (**Fig. 2L**). Since each distinct TCR clonal expansion can be regarded as a result of T cell proliferation and activation in response to the specific recognition of tumor antigen fragments, higher productive clonality in *Vhl-KO* MC38 tumors suggests the activation of different T cell clonotypes that are likely to target higher number of different tumor-specific antigens. Moreover, the max productive frequency measuring the frequency of dominant TCR clones also trended higher in *Vhl-KO* MC38 clones, despite not reaching statistical significance (**Fig. 2M**). Finally, the predominant T cell clones in *Vhl-KO* tumors had different TCR sequences than those in the control tumors (**Fig. 2N**), suggesting activation of different T cell subsets targeting different tumor antigens between *VHL*-WT and *Vhl*-KO tumors.

To evaluate the relative contribution of different immune effector subsets involved in antitumor immunity, we used antibodies to deplete CD4^+^ T cells, CD8^+^ T cells, and NK cells in mice bearing control and *Vhl-KO* MC38 tumors and observed their growth kinetics. Administration of an αCD8 antibody significantly accelerated the growth of VC and *Vhl-KO* tumors and completely abrogated the substantial growth delay of *Vhl*-KO tumors (**Fig. 2O-P**). On the other hand, administration of αCD4 and αNK1.1 antibodies did not affect tumor growth compared to those receiving isotype controls (**Fig. S2B-S2E**). Therefore, our data suggest a pivotal role for CD8^+^ T cells in mediating the growth delay in tumors with *Vhl* gene loss.

### VHL deficiency causes constitutive activation of type I interferons

To unravel the molecular mechanism of how *VHL* loss stimulates the antitumor immune response, we analyzed a publicly available dataset (GSE108229) comparing the transcriptional differences between parental 786-O RCC cells that carry a homozygous nonsense mutation in the *VHL* gene and 786-O cells with ectopic human *VHL (hVHL)* expression ^37^. Consistent with our findings from *in vivo* tumor growth delay studies, gene ontology (GO) analysis showed that *VHL* deficiency led to the activation of multiple immune-stimulating pathways (**Fig. 3A**). Notably, we found that IFNα/ý signaling is highly upregulated in *VHL*-deficient 786-O cells (**Fig. 3A**). To further validate this finding, we generated *VHL-KO* Caki-1 cells, a human clear cell carcinoma cell line with wild-type *VHL* expression (**Fig. S3A-S3B**). We then carried out RNAseq to identify transcriptomic changes in *VHL-KO* Caki-1 cells. Principal component analysis (PCA) of the RNAseq data demonstrated significant changes between VC and *VHL-KO* Caki-1 cells (**Fig. S3C**). Furthermore, gene set enrichment analysis (GSEA) identified significant enrichment of gene signatures involving IFNα/ý signaling and positive regulation of T cell mediated cytotoxicity in *VHL-KO* Caki-1 cells (**Fig. 3B, Fig. S3D**), consistent with our findings in 786-O cells and intratumoral lymphocytes infiltration in MC38 tumors. IFNα/β are type I interferons (IFNs) that can upregulate antigen presentation on the surface of tumor cells, which in turn stimulates the cross-presentation of tumor-specific antigens by professional antigen presentation cells (APCs) such as macrophages and dendritic cells, which are essential for antitumor immunity ^38^. Indeed, flow cytometry analysis showed a higher level of mouse H2K^b^/H2D^b^ expression on the surface of *Vhl-KO* MC38 cells compared to control MC38 cells (**Fig. S3D-S3E**). Furthermore, the mRNA levels of multiple human MHC class I (HLA) proteins were also substantially higher in *VHL-KO* Caki-1 cells (**Fig. S3F**). These data thus provide strong evidence indicating a vital role for *VHL* deficiency in stimulating the type I IFNs.

**Figure 3.**
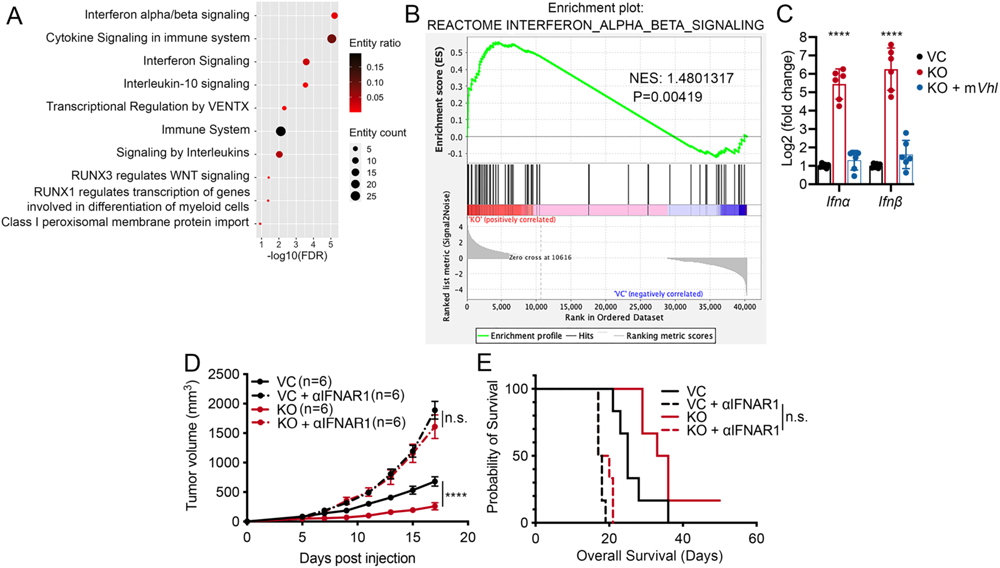
Type I interferon production is responsible for *VHL*-KO induced tumor growth suppression. **(A)** Gene ontology (GO) analysis shows top-10 pathways enriched in 786-O cells compared to 786-O cells with restored wild-type human *VHL (hVHL)* expression using publicly available data (GSE108229). **(B)** GSEA score curve plot showing the IFNα/ý signaling pathway enriched in *VHL*-KO Caki-1 cells. NES, normalized enrichment score. **(C)** Q-RT-PCR analysis of mRNA levels of *Ifnα* and *Ifný* in VC, *Vhl*-KO MC38 cells, and *Vhl*-KO MC38 cells with an exogenously expressed wild-type mouse *VHL (mVhl)* gene. We performed three independent experiments. Error bars represent mean ± SD. **(D-E)** Tumor growth and Kaplan-Meier survival analysis of C57BL/6J mice bearing VC and *Vhl*-KO MC38 tumors (n=6) following αIFNAR1 antibody treatments to suppress type I interferon signaling. About 100 μg isotype or αIFNAR1 antibodies per mouse were administered i.p. on days 5, 8, and 11 post tumor cell inoculation. Error bars represent mean ± SEM. P<0.05 is considered statistically significant. *P<0.05; **P<0.01; ***P<0.001; ****P<0.0001; n.s. not significant.

To further characterize the involvement of IFNα/ý signaling in suppressing the growth of *VHL-KO* murine tumors, we examined the expression of *Ifnα* and *Ifný* in VC and *Vhl-KO* MC38 cells, and *Vhl-KO* MC38 cells with restored *mVhl* gene expression. *Ifnα/ý* mRNA levels increased by approximately 32 folds and 64 folds in *Vhl-KO* MC38 cells, respectively. Notably, expression levels of both *Ifnα/β* reverted to control levels with the restoration of *mVhl* expression in *Vhl-KO* MC38 cells (**Fig. 3C**). To determine the functional importance of IFNα/β signaling in regulating antitumor immunity *in vivo*, we conducted a tumor growth delay experiment where we blocked the type I IFN signaling *in vivo* by injecting an αIFNAR1 antibody, which blocks the receptor for both IFNα/ý. Strikingly, blocking IFNAR signaling abrogated the tumor growth delay in *Vhl-KO* MC38 cells. Moreover, tumor growth from *Vhl-KO* MC38 tumors in mice receiving the αIFNAR1 antibody was even faster than that of control tumors, similar to VC MC38 tumors treated with the αIFNAR1 antibody (**Fig. 3D-E**). These results suggest that type I IFN signaling played an essential role in *Vhl-KO*-mediated antitumor immunity.

### Constitutive activation of the cGAS-STING pathway responsible for VHL loss-induced type I interferon activation

To determine the mechanism of type I IFN induction in *VHL-KO* cells, we further examined RNAseq data of *VHL-KO* Caki-1 cells using GSEA analysis. Our analysis demonstrated that *VHL* deficiency induced the expression of genes associated with the cyclic GMP-AMP synthase (cGAS)-stimulator of interferon genes (STING) pathway involved in sensing the cytoplasmic presence of double-stranded DNA (dsDNA) (Fig. 4A-B). The cGAS-STING pathway activation can induce interferon-stimulated genes (ISGs) and type I IFN production ^39–41^. Based on this analysis, we conducted immunoblot analysis of proteins associated with the VHL protein and the cGAS-STING pathway. As expected, we found that *VHL* loss in MC38 and Caki-1 cells caused the accumulation of HIFαs (**Fig. 4C-D**). In addition, it also upregulated the protein expression of STING and cGAS, and the phosphorylated form of the former in both cell lines (**Fig. 4C-D**), consistent with our RNAseq analysis. Furthermore, *VHL* loss activated several downstream effectors of the cGAS-STING axis, including the TANK binding kinase 1 (TBK1), interferon regulatory factor 3 (IRF3), and their phosphorylated forms, consistent with the activation of the cGAS-STING pathway (**Fig. 4C-D**). Furthermore, the observation of *VHL* gene loss-induced TBK1 activation was consistent with a recently published study ^42^. In addition, we also observed similar relationship between *Vhl*-KO and activation of the cGAS-STING pathway in the B16F10 melanoma cells (**Fig. S4A**). Moreover, to further establish the relationship between *VHL* loss and activation of the cGAS-STING axis, we re-expressed *mVhl* and *hVHL* in *Vhl-KO* mouse MC38 cells and *VHL-deficient* human 786-O cells, respectively (**Fig. 4C, 4E**). Immunoblot analysis showed that restoration of *VHL* abrogated the activation of STING, TBK1, and IRF3 in *VHL*-deficient cells (**Fig. 4C, 4E**).

**Figure 4.**
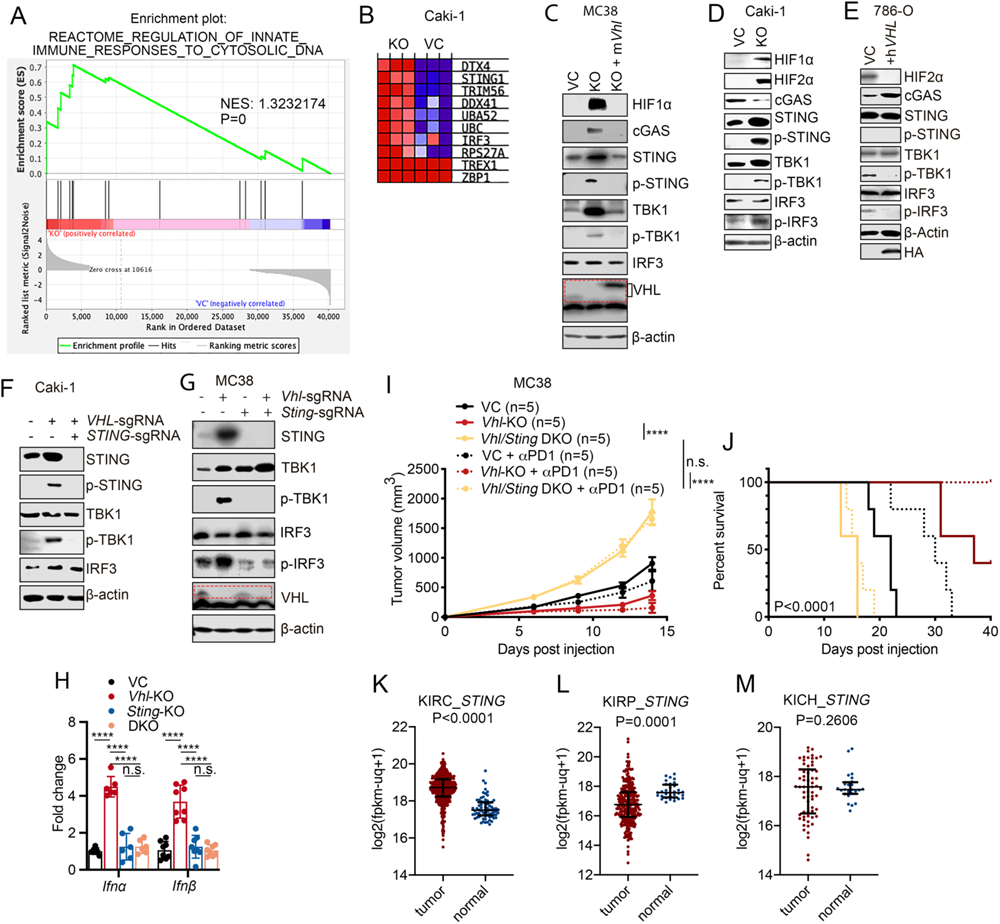
Activation of cGAS-STING signaling in *VHL*-KO cells and ccRCC human tumor tissues. **(A)** GSEA analysis of control and *VHL*-KO Caki-1 cells indicating elevated expression of genes involved in sensing cytosolic DNA. **(B)** Heatmap of top-10 differently expressed genes involved in regulating innate immune response to cytosolic DNA in control and *VHL*-KO Caki-1 cells. **(C-D)** Western blot analysis examining the effect of *Vhl/VHL* gene loss on cGAS-STING signaling and its downstream effectors in MC38 cells (C) and Caki-1 cells (D). In (C), ectopic mouse *Vhl* (m*Vhl*) was re-expressed in *Vhl*-KO MC38 cells. **(E)** Western blot analysis evaluating the effect of the ectopic expression of h*VHL* on cGAS-STING signaling in *VHL*-deficient 786-O cells. **(F-G)** Immunoblot analysis of that STING and TBK1 activation in *VHL*/*STING* DKO Caki-1 (F) and MC38 (G) cells. **(H)** Quantitative PCR determination of *Ifnα/β* levels in *Vhl*-KO, *Sting*-KO, and *Vhl*/*Sting* DKO MC38 cells. **(I-J)** Tumor growth (I) and Kaplan-Meier analysis (J) of C57BL/6 mice subcutaneously inoculated with VC, *Vhl*-KO, or *Vhl/Sting* DKO MC38 cells and treated with isotype control or αPD-1 antibody (100 μg per mouse) on day 6, 9, and 12 days post tumor cell inoculation. n=5 per group. **(K-M)** Comparisons of *STING* RNA expression levels between tumor tissues and adjacent normal tissues from TCGA_KIRC (clear cell renal cell carcinoma) (K), TCGA_KIRP (kidney renal papillary cell carcinoma) (L), and TCGA_KICH (kidney chromophobe renal cell carcinoma) (M) cohorts. *STING* expression is only significantly higher in ccRCC (KIRC) vs adjacent tissues where VHL is mutated in most tumors. Error bars represents mean ± SD. P<0.05 is considered statistically significant. *P<0.05; **P<0.01; ***P<0.001; ****P<0.0001; n.s. not significant.

To determine if the cGAS-STING pathway is essential for *VHL* gene loss-induced IFNα/ý production, we generated *VHL/STING* double-knockout (*DKO*) cells for Caki-1 and MC38 cells, respectively (**Fig. 4F-G**). Immunoblot analysis showed *STING-KO* abrogated *VHL* loss-induced type I interferon response in Caki-1 (**Fig. 4F**) and MC38 (**Fig. 4G**) cells. In contrast, genetic knockout of *Mda5,* a factor involved in sensing the cytoplasmic presence of double-stranded RNA, did not affect *Vhl-KO*-induced STING and TBK1 activation (**Fig. S4B**). In addition, *Sting-KO* eliminated *Vhl-KO*-dependent *Ifnα* and *Ifný* mRNA upregulation (**Fig. 4H**), suggesting the cGAS-STING-TBK1 axis is functionally responsible for *VHL* loss-induced type I IFN response. Notably, *Vhl/Sting* DKO completely reversed tumor growth delay and sensitization to anti-PD1 therapy caused by *Vhl* loss (**Fig. 4I-J**). Furthermore, the increased intratumoral lymphocyte infiltration observed in *Vhl*-KO tumors, especially those of CD3^+^T, CD8α^+^*a*, and CD8^+^GZMB^+^T cells, was significantly reduced in *Vhl/Sting* DKO tumors (**Fig. S4C-S4H**), further supporting the key role of cGAS-STING in *Vhl*-deficient tumor sensitivity to anti-PD1 blockades.

We also determined *STING* mRNA expression levels in three human TCGA renal cell carcinoma cohorts: KIRC, KIRP, and KICH. Compared with *STING* expression levels in adjacent normal tissues, only KIRC, representing clear cell renal cell carcinoma (ccRCC), showed significantly higher expression levels (**Fig. 4K**). In contrast, KIRP (kidney renal papillary cell carcinoma) (**Fig. 4L**) showed lower expression levels than normal tissues, while KICH (kidney chromophobe renal cell carcinoma) showed no difference (**Fig. 4M**). Because among three types of RCC, VHL function is lost in most KIRC but not in KICH or KIRP, the human RCC *STING* expression data were thus consistent with our preclinical observation.

### Mitochondrial dysfunction and cytoplasmic leakage of mtDNA responsible for VHL **deficiency-induced cGAS-STING activation**

We next attempted to determine the source of the cytosolic dsDNA that triggers cGAS-STING activation. We first fractionated VC and *VHL-KO* Caki-1 cellular lysates into pellet and cytosolic fractions and verified the success of the fractionation by using HSP60 and HDAC1 as markers for mitochondrial and nuclear fractions, respectively ^43–45^ (**Fig. 5A**). We then purified DNA from the cytosolic fractions and performed quantitative PCR analysis using primers that amplify nuclear (nucDNA) and mitochondrial (mtDNA) encoded genes. Our results indicate no significant differences in the amount of cytosolic nucDNA between VC and *VHL-KO* Caki-1 cells (**Fig. 5B**). In contrast, we consistently detected more mtDNA in the cytosolic fraction of *VHL-KO* than VC Caki-1 cells (**Fig. 5B**), strongly suggesting mtDNA leakage into the cytoplasm as the trigger for cGAS-STING activation in *VHL-KO* cells. We also made similar observations in murine MC38 and B16F10 tumor cells (**Fig. S5A-S5D**). We next attempted to detect the cytoplasmic presence of dsDNA by immunofluorescence staining in VC and *VHL-KO* Caki-1 cells. Using an antibody-specific for dsDNA, we observed that most cytoplasmic DNA was co-localized with HSP60 in the mitochondria in control cells (**Fig. 5C**). However, a substantial amount of dsDNA did not co-localize with HSP60 in *VHL-KO* cells (**Fig. 5C, Fig. S5E**), suggesting extra-mitochondrial locations for the dsDNA. These data thus provide correlative evidence that mtDNA leaked into the cytoplasm in *VHL-KO* cells is responsible for the observed cGAS-STING activation. To determine if mtDNA is functionally required for cGAS-STING activation, we used an established protocol to deplete mtDNA by ethidium bromide (EtBr) ^46–48^. After a 21-day EtBr treatment, we could deplete most cytosolic mtDNA as confirmed by immunostaining using an anti-dsDNA antibody in Caki-1(**Fig. 5D**), and MC38 (**Fig. S5F**) cells. Our data suggest that EtBr treatment substantially suppressed cGAS-STING and/or type I IFN response *VHL-KO* Caki-1 (**Fig. 5E**) and MC38 (**Fig. 5F, Fig. S5G**) cells. These results, therefore, support the notion that mtDNA leakage into the cytoplasm by *VHL-KO* is responsible for activating the cGAS-STING pathway and inducing type I IFN production.

**Figure 5.**
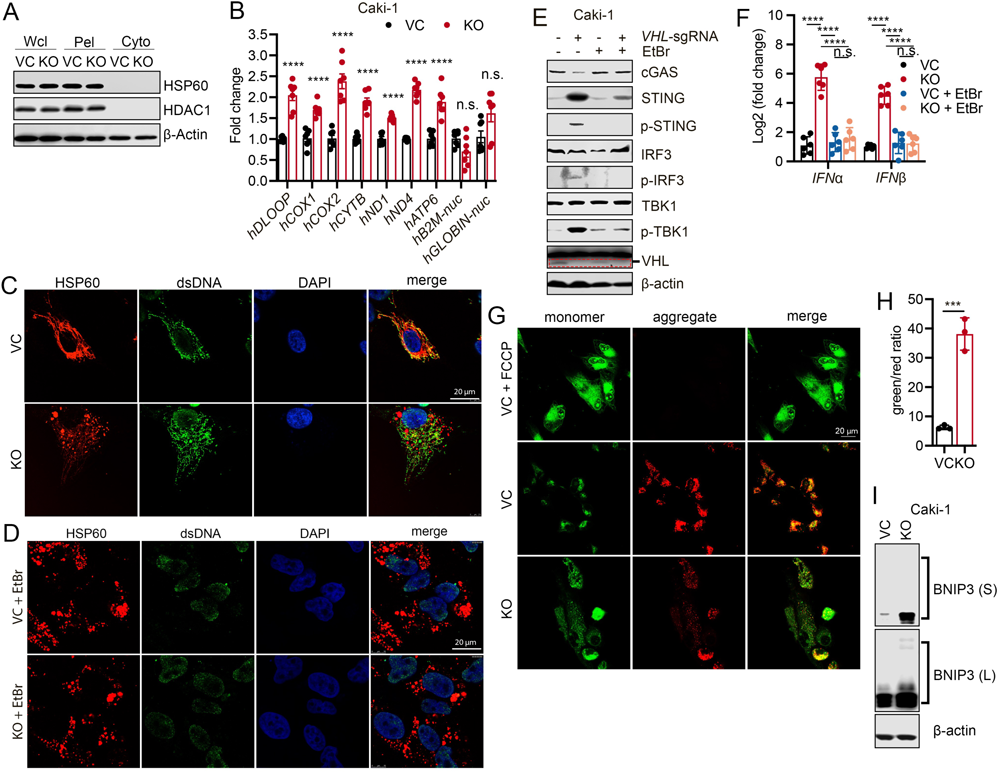
Cytoplasmic leakage of mitochondrial DNA as the key driver of cGAS-STING activation in *VHL*-deficient cancer cells. **(A)** Immunoblot analysis validation of the absence of nuclear (*HADC1* as the marker) and mitochondrial (*HSP60* as the marker) proteins. Wcl, whole cell lysate; Pel, pellet fraction; Cyto, cytosolic fraction. **(B)** Q-RT-PCR measurement of mtDNAs in the cytosols of VC and *VHL*-KO Caki-1 cells (n=3). Error bars represent mean ± SEM. **(C)** Immunofluorescent localization of the mitochondria (as detected by HSP60) and dsDNA in the cytoplasm of VC and *VHL*-KO Caki-1 cells. Scale bar, 20μm. **(D)** Immunofluorescence detection of cytoplasmic dsDNA in VC and *VHL*-KO Caki-1 cells after a 21-day treatment in 100ng/ml EtBr to deplete mtDNA. Scale bar, 20μm. **(E)** Immunoblot analysis of proteins involved in cGAS-SITNG signaling in VC and *VHL*-KO Caki-1 cells treated with vehicle or 100ng/ml EtBr for 21 days. **(F)** Q-RT-PCR analysis of the mRNA levels of *IFNα* and *IFNϕ3* in VC and *VHL*-KO Caki-1 cells treated with vehicle or 100ng/ml EtBr for 21 days (n=6). Error bars represent mean ± SD. **(G-H)** Mitochondrial membrane potential (MtMP) characterization by JC-1 staining in VC and *VHL*-KO Caki-1 cells as detected by immunofluorescence microscopy (G) and flow cytometry (H). Scale bar, 20μM. Error bars represent mean ± SD. *P<0.05; **P<0.01; ***P<0.001; ****P<0.0001; n.s. not significant**. (I)** Immunoblot analysis of BNIP3 expression in VC and *VHL*-KO Caki-1 cells.

Since mtDNA leakage into the cytoplasm is most likely due to mitochondria damage, we sought to understand how *VHL* deficiency may disrupt mitochondrial functions. MtDNA leakage is usually associated with lowered mitochondrial membrane potential (MtMP), closely associated with the permeability changes in mitochondrial membranes ^49^. To determine whether *VHL* deficiency influences mitochondrial membrane permeability, we assessed mitochondrial permeability by using the JC-1 fluorescent probe ^50,51^ in VC and *VHL-KO* cells. Flow cytometry analysis found a significantly higher green(monomer) / red(aggregate) fluorescence ratio of JC-1 in *VHL-KO* cells, indicating significantly lowered MtMP in VHL-deficient Caki-1 cells (**Fig. 5G-H, Fig. S6A**) and MC38 cells (**Fig. S6B-S6D**). These data suggest that *VHL* deficiency causes a significant reduction in MtMP. How does *VHL* deficiency cause a decrease in MtMP? Previous studies have shown that hypoxia can induce the expression of BNIP3^52,53^, which can increase mitochondrial outer membrane permeability and decrease membrane potential^54,55^. Therefore, we hypothesized that *VHL* deletion was akin to subjecting cells to a condition of perpetual hypoxia by permanently upregulating HIFα expression, which could upregulate BNIP3 expression. Indeed, our data indicate that *VHL* knockout significantly increased the level of BNIP3 (**Fig. 5I**), consistent with our hypothesis.

### Increased HIF1/2α and BNIP3 levels in VHL-deficient cells are responsible for mitochondrial dysfunction and mtDNA leakage

We next sought to understand the molecular mechanism of how *VHL* deficiency causes the reduction in mitochondrial membrane potential. Because HIF1α and HIF2α are direct targets of VHL and are associated with hypoxia-induced mitochondrial dysfunction ^2,3,56–58^ and *VHL*-deficient cells had elevated levels of HIF1α and HIF2α (**Fig. 4C-D**), we decided to test whether forced expression of exogenous *HIF1α* and *HIF2α* could activate the cGAS-STING pathway. Therefore, we introduced into Caki-1 cells mutant versions of h*HIF1α* or h*HIF2α*, both of which are resistant to prolyl hydroxylase (PHD)-induced hydroxylation and thus resistant to VHL-mediated degradation. Ectopic expression of mut*HIF1α* significantly enhanced monomer/aggregates ratio in JC-1 stained Caki-1 cells compared with control cells (**Fig. 6A, Fig. S6E**), indicating a dramatic reduction in MtMP. In comparison, expression of mut*HIF2α* only moderately reduced MtMP (**Fig. 6A, Fig. S6E**). Furthermore, expression of mut*HIF1α*, but not mut*HIF2α*, induced elevated expression of STING and activation of TBK1 (as indicated by detection of p-TBK1) in Caki-1 cells (**Fig. 6B**), similar to those observed in *VHL*-KO cells (**Fig. 4D**). Furthermore, similar to *VHL* knockout, mut*HIF1α* also induced the activation of BNIP3 in Caki-1 cells (**Fig. 6B**), consistent with it being a direct transcriptional target of HIF1α. Our data, therefore, indicate that *HIF1α* overexpression is sufficient to activate type I IFN response.

**Figure 6.**
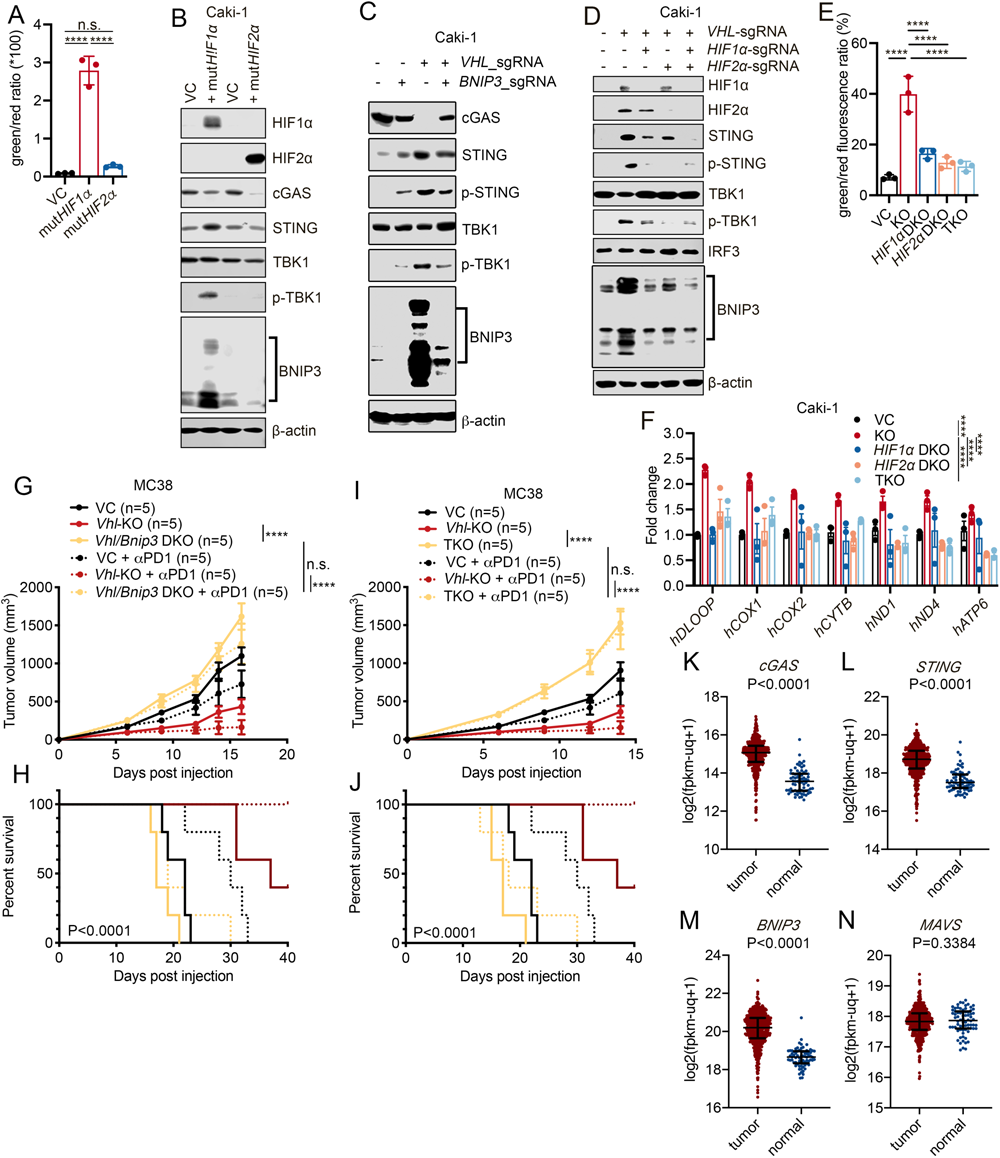
Elevated HIF1α and HIF2α levels are essential and sufficient for *VHL*-deficiency induced mitochondrial dysfunction and cGAS-STING activation. **(A)** Quantitative JC-1-based MtMP measurements in Caki-1 cells expressing VC, degradation-resistant mut*HIF1α* (h*HIF1α*-p402A/p564A), or mut*HIF2α* (h*HIF2α*-p405A/p531A) by flow cytometry. Error bars represent mean ± SD. **(B)** Immunoblot analysis of cGAS-STING signaling proteins and BNIP3 in Caki-1 cells expressing exogenous VC, mut*HIF1α*, or mut*HIF2α* genes. **(C)** Immunoblot analysis of cGAS-STING signaling proteins in VC, *BNIP3*-KO, *VHL*-KO, and *VHL*/*BNIP3* DKO Caki-1 cells. **(D)** Immunoblot analysis of cGAS-STING signaling proteins and BNIP3 in VC, *VHL*-KO, *VHL*/*HIF1α* DKO, *VHL*/*HIF2α* DKO, and *VHL*/*HIF1α*/*HIF2α* TKO Caki-1 cells. **(E)** JC-1-based MtMP measurements in VC, *VHL*-KO, *VHL*/*HIF1α* DKO, *VHL*/*HIF2α* DKO, and *VHL*/*HIF1α*/*HIF2α* TKO Caki-1 cells by flow cytometry. Error bars represent mean ± SD. **(F)** Quantitative PCR measurements of cytosolic mtDNAs in VC, *VHL*-KO, *VHL*/*HIF1α* DKO, *VHL*/*HIF2α* DKO, and *VHL*/*HIF1α*/*HIF2α* TKO Caki-1 cells. **(G-H)** Tumor growth (G) and Kaplan-Meier analysis (H) of C57BL/6 mice subcutaneously inoculated with VC, *Vhl*-KO, or *Vhl/Bnip3* DKO MC38 cells and treated with isotype control or αPD-1 antibody (100 μg per mouse). **(I-J)** Tumor growth (I) and Kaplan-Meier analysis (J) of C57BL/6 mice implanted with VC, *Vhl*-KO, or *Vhl/Hif1α/Hif2α* TKO MC38 cells and treated with isotype control or αPD-1 antibody (100 μg per mouse). G and I, treatments were given on day 6, 9, and 12 days post tumor cell inoculation. n=5 per group. **(K-N)** Normalized mRNA expression levels of *cGAS* (K), *STING* (L), *BNIP3* (M), and *MAVS* (N) in tumor tissues and normal tissues from the TCGA_KIRC(ccRCC) cohort. Tumor tissues have higher levels expressions with the exception of MAVS, which is not part of cGAS-STING dsDNA-sensing pathway. Shown are results from three independent experiments. Error bars represent mean ± SEM. Two-way ANOVA. *P<0.05; **P<0.01; ***P<0.001; ****P<0.0001; n.s. not significant.

To test whether BNIP3, that is upregulated by *VHL*-KO-mediated HIF1α accumulation, is required to promote cGAS-STING signaling activation, we further generated a mixed cell population of *BNIP3*-KD and *VHL/BNIP3* DKD using CRISPR-Cas9 system (**Fig. 6C, Fig. S6F**). *VHL/BNIP3* DKD abrogated *VHL*-KO-induced activation of STING and TBK1, the levels of which were comparable in *VHL/BNIP3* DKD and *BNIP3*-KD alone cells (**Fig. 6C, Fig. S6F**). Therefore, elevated cGAS-STING signaling due to *VHL* loss is most likely regulated by VHL-HIF1α-BNIP3 axis.

We further investigated whether HIF1α and HIF2α were essential for reducing MtMP and causing cytosolic mtDNA leakage in VHL-deficient cells. For this purpose, we generated *VHL/HIF1α DKO, VHL/HIF2α DKO*, and *VHL/HIF1α/HIF2α* triple knockout (TKO) Caki-1 cells, respectively (**Fig. 6D**). Notably, knockout of either *HIF1α* or *HIF2α* or both significantly reduced *VHL*-KO-induced activation of BNIP3 and cGAS-STING signaling (**Fig. 6D**). Therefore, both HIF1α and HIF2α stabilization can contribute to the activation of cGAS-STING signaling. In agreement with this finding, *VHL-*KO-mediated reduction of mitochondrial membrane potential was largely rescued in *VHL/HIF1α* DKO*, VHL/HIF2α* DKO, and *VHL/HIF1α/HIF2α* TKO cells, suggesting that both HIF1α and HIF2α are required for the *VHL* deficiency-induced mitochondrial membrane permeabilization (**Fig. 6E, Fig. S6G**). Consistent with this finding, individual or combined *HIF1/2α* deletions in *VHL-KO* cells significantly reduced cytosolic mtDNA (**Fig. 6F**), while changes of cytosolic nuclear DNA were insignificant (**Fig. S6H**). Importantly, further knockout of *Bnip3* and *Hif1α/2α* abrogated tumor growth delay observed in *Vhl-KO* MC38 tumors. In addition, loss of *Bnip3* or *Hif1α/Hif2α* in *Vhl*-deficient MC38 tumors also attenuated their sensitivity to ICB treatments significantly (**Fig. 6G-J**). Taken together, our results suggest that HIF1α and HIF2α are the critical effectors downstream of VHL responsible for reducing mitochondrial membrane potential and increasing membrane permeability via BNIP3 to activate cGAS-STING signaling.

Many human tumors are resistant to immunotherapy because they downregulate STING signaling by various mechanisms, ^59,60^ a fact that testifies to the importance of the cGAS-STING pathway in mediating tumor response to ICB therapy. In order to see if the cGAS-STING pathway is indeed upregulated in ccRCC as has been shown in our experiments, we analyzed the mRNA expression levels of *cGAS* and *STING* in a cohort of ccRCC patients from the TCGA PanCancer cohort. Our analysis indicated that both *cGAS* (**Fig. 6K**) and *STING* (**Fig. 6L**) were expressed at higher levels in tumor vs adjacent normal tissues, thereby confirming the activation of the cGAS-STING pathway. Furthermore, *BNIP3* was similarly expressed at higher levels in tumor tissues (**Fig. 6M**). Therefore, data from ccRCC patients are consistent with activation of the mtDNA-cGAS-STING pathway. In contrast, expression of MAVS, a vital component of the dsRNA-sensing pathway, showed no difference between tumor and adjacent normal tissues (**Fig. 6N**). Taken together, our analysis of the TCGA human ccRCC data is consistent with the activation of mtDNA-cGAS-STING pathway in these patients, the majority of whom have VHL deficiencies.

### VHL deficiency leads to better response to ICB therapy in human malignancies

To study the influence of *VHL* loss on the treatment outcome of ccRCC patients undergoing ICB therapy, we integrated three publicly available cohorts of ICB-treated ccRCC patients study the influence of *VHL* loss on the treatment outcome of ccRCC patients undergoing ICB therapy, we integrated three publicly available cohorts of ICB-treated ccRCC patients ^26,28,61^. These patients received ICB monotherapy without additional combined treatments, including but not limited to mTOR inhibitors, tyrosine kinase inhibitors, and VEGF/VEGFR blockades. First, we analyzed the overall survival (OS) of patients with wild-type and mutant *VHL* (**Fig. 7A**). We found that patients with *VHL* mutations had significantly better OS (**Fig. 7A**) compared to those with wild-type *VHL*, while mutations in other genes, including *PBRM1, SETD2 BAP1*, and *KDM5C* that were frequently mutated in ccRCC, did not influence ICB treatment (**Fig. S7A-S7D**). These data suggest that *VHL* deficiency is uniquely associated with susceptibility to ICB therapy in the ccRCC among the most frequently mutated genes. In contrast, the analysis of non-ICB treated ccRCC patients from Pan-Cancer cohorts on cBioPortal (http://www.cbioportal.org) indicates similar OS between patients with wild-type *VHL* and those with *VHL* mutations (**Fig. S7E**), suggesting that VHL mutations do not play a role in non-ICB treatment.

**Figure 7.**
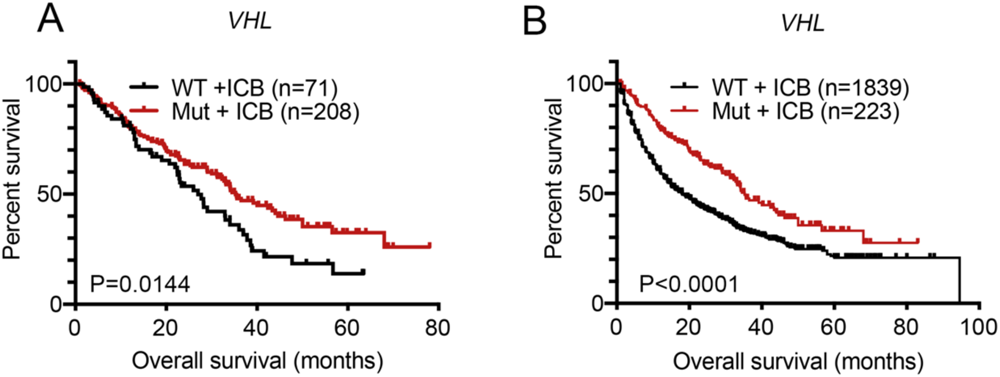
*VHL* loss predicts for better human tumor response to ICB therapy. **(A)** Kaplan-Meier survival analysis of ICB-treated ccRCC patients with WT *VHL* or *VHL* gene mutations. **(B)** Kaplan-Meier survival analysis of patients with WT *VHL* or *VHL* gene mutations undergoing ICB therapy across multiple cancers types.

We next analyzed the OS of those patients with *VHL* gene mutations against those with wild-type *VHL* genes in ten ICB-treated patient cohorts across different cancer types, including eight from cBioportal immunogenic studies and two from CheckMate cohorts ^26,28,61–67^. In this multi-cancer cohort, the vast majority of the patients have non-RCC malignancies, including lung cancer, melanoma, bladder cancer, colon cancer, etc. We found that those with *VHL* gene mutations again showed a significant survival advantage over those with wild-type *VHL* genes (**Fig. 7B**). Therefore, our data suggest that *VHL* mutations are associated with the responsiveness to ICB therapy in different cancer types.

## Discussion

Our finding of *VHL* gene loss causing constitutive activation of the cGAS-STING signaling and increased intratumoral lymphocyte infiltration provides critical insights into how ccRCC unexpectedly responds well to ICB therapy ^18,21^ despite its very moderate mutation burden and relative lack of neoantigens^24^. These insights are critical in understanding the biology of ccRCC and its treatment, given that *VHL* gene mutations are the dominant genetic event in ccRCC^14^, occurring in as many as 91% of all cases^7^. The mechanistic insights gained from our study thus provide a solid foundation to understand ccRCC responsiveness to ICB therapy (**Fig. 8**).

**Figure 8:**
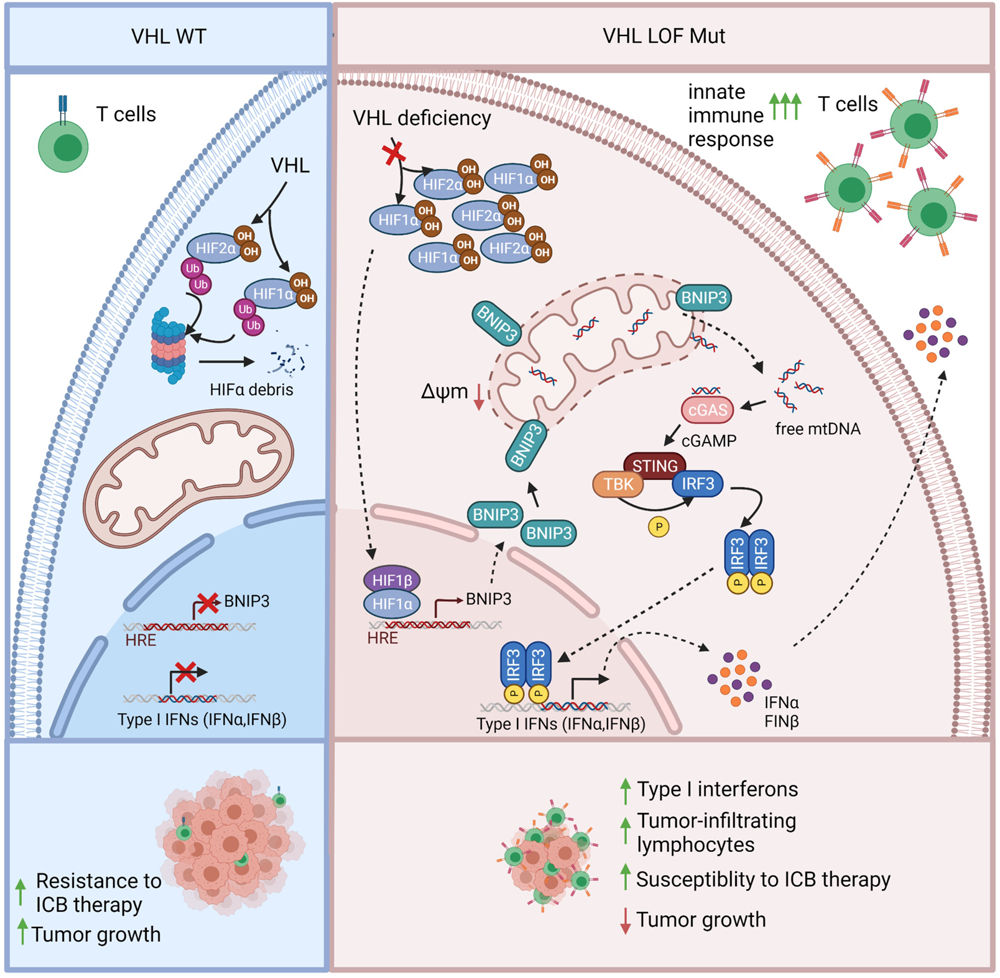
Schematic diagram elucidating the regulatory mechanisms of VHL loss in enhancing tumor susceptibility to immunotherapy.

How do we reconcile the paradoxical roles of mutations in the *VHL* gene, which promote the development of ccRCC, also predispose ccRCCs to better response to ICB therapy. The paradox may have to do with the dual roles of cGAS-TING pathway, which acts downstream of VHL. Indeed, chronic cGAS-STING has been reported to promote tumor invasion and metastasis^68^. Therefore, it is very likely that *VHL* mutations promote ccRCC development through cGAS-STING activation. Consistent with our finding of *VHL* deficiency-induced cGAS-STING activation, TBK1, a key factor downstream of cGAS-STING, has recently been identified as a synthetic lethal target because of its constitutive activation in ccRCC with *VHL* loss^42^. Furthermore, the paradoxical roles of *VHL* mutations in promoting tumor development while making them susceptible to ICB therapy are not unique. Similar examples include mutations in the ATM gene and mismatch repair genes. Both are well-established tumor suppressor genes whose mutations promote tumorigenesis but make them more susceptible to immunotherapy^46,69^.

Unlike our findings, many published clinical studies did not show an improved response to ICB treatment in *VHL*-mut ccRCC patients. We do not know the underlying mechanism for this discrepancy. However, we surmise there could be several factors. One confounding factor might be that patients from most of the published studies also received different co-treatments. For example, in Javelin Renal 101^70^, ImMotion150^34^, and ImMotion 151^71^ RCC cohorts, patients receiving anti-PD-L1 treatment were also treated with VEGF inhibitors or anti-VEGF treatments concurrently, while the ccRCC patients we analyzed (**Fig. 7A**) (including 123 patients from CheckMate 009/010/025^61^, 35 patients from DCFI^28^, and 121 patients from MSKCC^26^) did not undergo such co-treatments. The other factor undermining the use of *VHL* genomic mutation as a biomarker for ICB therapy may be that *VHL* genomic mutations may not fully reflect the actual functional status of the *VHL* gene. In fact, even those ccRCC patients with no *VHL* genomic mutations may have compromised VHL functions. For example, methylation-mediated *VHL* gene inactivation or loss of heterozygosity (LOH), which may occur in up to 98% of ccRCC patients^37,72^, may also disrupt VHL function. Therefore, even though we believe that compromised VHL function underlies the overall ccRCC responsiveness to ICB therapy, genomic mutations may not be a suitable biomarker for predicting individual ccRCC response to ICB therapy because of the wide prevalence of compromised VHL functions in ccRCC.

In addition, even though *VHL* mutations occur primarily in ccRCC, an examination of the Cancer Genome Atlas (TCGA) database ^73^ indicates that many non-ccRCC malignancies do possess *VHL* mutations, which can occur in breast cancer, colon cancer, lung cancer, and melanoma at much lower frequencies (**Table S1-S3**). Under these circumstances, it is conceivable to use *VHL* gene loss as a biomarker for ICB therapy, similar to mismatch repair deficiency, which occurs rarely but is FDA-approved for use as a tumor type-agnostic biomarker for ICB therapy ^69,74^. However, further studies are needed to evaluate such a tantalizing possibility. On the other hand, the frequencies of *VHL* mutations in non-ccRCC malignancies are exceedingly low when compared with ccRCC. We do not fully understand why *VHL* mutations are much less common in other malignancies. However, based on the present study, we speculate that such mutations may subject the host cells to higher levels of immune surveillance and elimination due to elevated levels of type I interferon production, significantly lowering their chances for malignant transformation.

In addition, the constitutive cGAS-STING activation in *VHL*-deficient tumors can potentially create additional therapeutic opportunities beyond ICB immunotherapy. For example, *VHL*-deficient tumors may be more susceptible to chimeric antigen receptor T (CAR-T) therapy due to elevated levels of type I interferons in the tumor microenvironment. Indeed, recent studies have shown that NKG2D CAR-engineered T cells can synergize with STING agonists in suppressing the growth of pancreatic tumors^75^. By the same token, adoptively transferred TILs (tumor-infiltrating lymphocytes) may have a better chance of penetrating the tumor mass and staying active due to a more hospitable tumor immune microenvironment created by constitutive cGAS-STING activation. Another possibility is using STING agonists in combination with ICB therapy in RCC. The agonists may “supercharge” the constitutively high STING levels and create a more immune-stimulating tumor microenvironment that further enhances ICB therapy.

In conclusion, our study reveals novel insights into the *VHL* deficiency-induced cGAS-STING activation, providing a mechanistic underpinning of why ccRCCs respond to ICB therapy despite its very moderate TMB. Finally, it also suggests potential novel treatment strategies for targeting ccRCC and other *VHL*-deficient tumors based on constitutive cGAS-STING and type I interferon activation.

## Materials and methods

### Cell Culture

We purchased the Renca mouse renal adenocarcinoma cells (ATCC® CRL2947™), B16F10 mouse melanoma cells (ATCC® CRL6475™), Caki-1 human clear cell carcinoma cells (ATCC® HTB-46™), and 786-O human renal cell adenocarcinoma (ATCC® CRL1932™) from the Cell Culture Facility of Duke University School of Medicine. In addition, we obtained the MC38 mouse colon adenocarcinoma cells from Kerafast (Boston, MA). All cells undergo periodic mycoplasma testing using the Universal Mycoplasma Detection Kit from ATCC (Cat #30-1012K) to ensure they are mycoplasma-free.

### Antibodies and Reagents

Trustain FcX^TM^ anti-mouse CD16/32 (Cat #101320), FITC anti–mouse CD45 (30-F11; Cat #103108), Pacific blue anti–mouse CD3 (145-2c11; Cat #100334), Alexa Fluor 647 anti–mouse CD4 (GK1.5; Cat #100424), APC/Fire^TM^ 750 anti– mouse CD8a (53-6.7; Cat #100766), phycoerythrin (PE) anti–mouse NK1.1 (PK136; Cat #108707), APC anti–γ/δTCR (GL3; Cat #118116), PE anti–mouse F4/80 (BM8; Cat #123110), phycoerythrin (PE) anti–human/mouse Granzyme B (QA16A02; Cat #372207), Alexa Fluor 647 anti–mouse IFN-γ (XMG1.2; Cat #505816), PE anti–mouse FOXP3 (MF-14; Cat #126404), and PE anti-mouse H2-Kb/H2Db antibody (28-8-6, cat#114607) were purchased from BioLegend. Anti-VHL (VHL40; Cat #sc-135657) was purchased from Santa Cruz Biotechnology. Anti-GAPDH (cat #60004-1-Ig) was purchased from Proteintech. Anti-cGAS (D3080; cat# 31659), anti-cGAS (D1D3G; Cat #15102), anti-STING (2P2F; Cat# 13647), anti-p-STING (Ser365) (D8F4W; Cat #72971), anti-p-STING (Ser366) (D7C3S; Cat #19781), anti-TBK1/Nak (D1B4, Cat #3504), anti–p-TBK1/p-Nak (Ser172) (D52C2, Cat #5483), anti-IRF-3 (D83B9; Cat #4302), anti-p-IRF3 (Ser396) (D601M; Cat #29047), anti-HA-Tag (C29F4; Cat #3724), anti-HSP60 (D307; Cat #4870), anti-HDAC1 (Cat #2062), anti-BNIP3 (D7U1T; Cat #44060), and anti-MDA5 (D74E4, Cat#5321) were purchased from Cell Signaling Technology. Anti-actin (ACTN05-C4; Cat #MA5-11869) was purchased from Thermo Fisher Scientific. Anti-HIF-1 alpha (H1alpha67; Cat #NB100-105) and anti-HIF-2 alpha/EPAS1 (Cat #NB100-122) were purchased from Novus Biologicals. Anti-dsDNA (AC-30-10; Cat #CBL186) was purchased from Millipore Sigma. Alexa Fluor^®^ 488 goat anti-mouse IgG (Cat #A28175) and Alexa Fluor^®^ 555 goat anti-rabbit IgG (Cat #A27039) were purchased from Invitrogen by Thermo Fisher Scientific.

### CRISPR/Cas9-mediated gene knockout

We used the CRISPR/Cas9 system to generate gene-specific knockout cells. We designed single guided RNAs (sgRNAs) using a public domain web-based CRISPR sgRNA design tool CHOPCHOP (https://chopchop.cbu.uib.no). Table S4 lists individual sgRNA sequences targeting different genes. When generating Human *VHL*-KO, human *STING*-KO, human *HIF1α-* KO, human *HIF2α-*KO, mouse *Sting*-KO, and mouse *Mda5*-KO human and murine cancer cells, we used the lentiCRISPRv2 vector (Addgene #52961) following a published protocol from the Zhang lab ^76^. We digested LentiCRISPRv2 with BsmBI (NEB; Cat #R0580) and gel purified it using the Gene JET Gel extraction kit (Thermo Fisher Scientific; Cat #K0692). Oligos encoding sgRNA sequences were phosphorylated, annealed, and subsequently ligated into digested LentiCRISPRv2.

To produce sgRNA-encoding lentivirus vectors, we used HEK293T cells co-transfected with lentiviral constructs encoding the target sgRNAs and second-generation packaging plasmids psPAX2 (Addgene #12260) and pMD2.G (Addgene #12259) following the instruction from the Trono Lab (https://www.epfl.ch/labs/tronolab/laboratory-of-virology-and-genetics/lentivectors-toolbox/).

To generate clonal knockout cell lines, we infected target cells with sgRNA-encoding CRISPR/Cas9 lentivirus and cultured them in a complete cell growth medium, selected with puromycin (1μg/ml for Caki-1) for 7-10 days. Cells were then collected to test the expression of target genes by immunoblot.

To generate clonal *VHL*-KO Caki-1 cells, cells were seeded in 96-well plates after a 7-day puromycin selection and screened for pure knockout clones verified by immunoblot analysis. To generate *VHL/STING* double knockout (DKO) cells, including *VHL/HIF1α* DKO, *VHL/HIF2α* DKO, or *VHL/HIF1α/HIF2α* triple knockout (TKO) Caki-1 cells, we cloned sgRNAs sequences encoding *HIF1α*, and/or *HIF2α* into digested lentiCRISPRv2 neo vector (Addgene #98292), respectively. We then infected *VHL*-KO Caki-1 cells with the sgRNA-encoding lentivirus and selected the cells with neomycin (2mg/ml) for 10-14 days. Lentivirus prepared from lentiCRISPRv2 neo vector was used to generate control cells. We then detected protein levels of STING, HIF1α, and/or HIF2α knockdown (KD) by western blot. To generate *VHL/STING* DKO and *VHL/MDA5* DKO Caki-1 and MC38 cells, we infected *VHL-KO* Caki-1 cells with sgRNA-encoding lentivirus and cultured them in complete DMEM medium, and selected with puromycin (5μg/ml for MC38 cells and 1μg/ml for Caki-1 cells) for 7 days. We then seeded the cells in 96-well plates for single clone selection. We then verified the cells’ *VHL/STING* DKO and *VHL/MDA5* DKO status by immunoblot analysis.

To generate Vhl knockout in Renca, B16, or MC38 cells, we digested the px330-mcherry (Addgene #98750) and pSpCas9(BB)-2A-GFP(PX458) (Addgene #48138) with BbsI (NEB; Cat #R0539S), and then gel-purified, and ligated the vectors with annealed oligos encoding sgRNA_1 or sgRNA_2 targeting *Vhl*. We then transfected the vectors in the murine tumor cells with sgRNA-encoding constructs using Lipofectamine^TM^ 2000 (Thermo Fisher Scientific; Cat #11668019). Cells transiently transfected with px330-mCherry and PX458 vectors alone were as controls (VC). We then cultured the cells using a complete DMEM medium [DMEM with 10% FBS and 100U/ml penicillin-Streptomycin (Gibco^TM^ by Life Technologies; Cat #15140122)] for 6 days and subjected them to FACS sorting (Duke Cancer Institute flow cytometry core facility). We then seeded GFP^+^ mCherry^+^ cells in 96 well plates for single clone selection. We screened for VHL-KO single clones by immunoblot analysis. Table S4 shows the sgRNA primer sequences used for targeting various genes in this study.

### Exogenous gene expression

We modified a commercially available LentiORF pLEX vector (Thermo Scientific; Cat #OHS4735) with EF-1α and used it to generate constructs for exogenous gene expression. Gene of interest was amplified by PCR using the Phusion^®^ High Fidelity DNA Polymerase. Specifically, we used HA-VHL-pRC/CMV (Addgene #19999) and HA-tagged mouse VHL ORF Clone (Origene; Cat #MR201630) as templates to amplify human and mouse VHL, respectively. In addition, we amplified mutant human HIF1/2α using the pcDNA3-HA-HIF1α(P402A/P564A) (Addgene #18955) plasmid and the pcDNA3-HA-HIF2α(P405A/P531A) plasmid (Addgene #18956). Modified pLEX vector was used for lentiviral production as controls. Table S5 lists the primer sequences for PCR reactions.

### Immunoblot analysis

To prepare cellular lysates, we washed the cells quickly with ice-cold 1X PBS buffer twice and immediately lysed them in 1× RIPA buffer (Sigma; Cat #R0278) supplemented with 1× protease inhibitors (Sigma; Cat #P8340) on ice using a cell scraper. We then collected the lysates in a 1.5 ml tube and incubated them on ice for 10 minutes, then centrifuged them at 15,000 rpm for 15 minutes at 4°C. We then transferred the supernatants to a new 1.5 ml tube and boiled them for 5 minutes. We then loaded equal amounts of lysates into SDS-PAGE gels. After electrophoresis, we transferred the proteins to PVDF membranes, incubated them with primary antibodies overnight at 4°C, and then incubated them with HRP-conjugated secondary antibodies at room temperature for 1 hour. Afterward, we incubated the membranes with the SuperSignal^TM^ West Pico PLUS Chemiluminescent Substrate (Thermo Fisher Scientific; Cat #34580) and detected target signal strength with the Odyssey^®^ Fc imaging system (LI-COR^®^ Biosciences).

### Tumor growth experiments in mice

We purchased six-week-old C57BL/6J and Balb/C female mice from the Jackson Laboratory. Duke University Institutional Animal Use and Care Committee (IACUC) approved all mouse experiments in this study. For *in vivo* tumor growth experiments, we inoculated about 1*10^6^ Renca cells, 1*10^5^ B16F10 cells, and 5*10^5^ MC38 cells (suspended in 50 μl 1× PBS) subcutaneously into the right flanks of syngeneic mice, respectively. We measured tumor sizes by measuring the longest (length) and shortest (width) dimensions of the tumors every 2-3 days using a digital caliper and calculated tumor volumes using the following formula: (length) × (width)^2^/2. We euthanized mice when their volumes reached 2000 mm^3^.

In conducting immunotherapy in mice, we injected the antibodies intraperitoneally (i.p.), either with 100 μg of rat IgG2 isotype control (clone 2A3; Bio X Cell; Cat #BE0089), or 100μg of rat anti-mouse αPD-1 antibody in 150 μl 1×PBS per mouse on Day 5, 8, and 11 for Renca cells, on Day 7, 10, 13, and 16 for B16F10 cells, and on Day 6, 9, 12 for MC38 cells after tumor inoculation, respectively.

In experiments involving αIFNAR1 antibody treatments, we injected mice (i.p.) bearing MC38 tumors with 200 μg mouse IgG1 isotype control (clone MOPC-21; BioXCell; Cat #EB0083) or 200 μg mouse αIFNAR1 antibody (clone MAR1-5A3; BioXCell; Cat #BE0241) in 150 μl PBS per mouse on Day 5, 8, and 11, respectively.

In studies involving *in vivo* lymphocyte depletion, we injected 100 μg αCD4 antibody (Clone GK1.5; BioXCell; Cat #BE0003-1), or αCD8b antibody (Clone 53-5.8; BioXCell; Cat #BE0223), or 100 μg αNK1.1 antibody (Clone PK136; BioXCell; Cat #BE0036) into each MC38 tumor-bearing mouse on Day 1, 4, and 7 to deplete CD4^+^ T cells, CD8^+^ T cells, and NK cells, respectively. We also administered an equal amount of IgG isotype antibodies to control mice.

### Flow cytometry

To analyze tumor-infiltrating lymphocytes, we implanted about 5*10^5^ vector control or *Vhl*-KO, or *Vhl/Sting* DKO MC38 cells subcutaneously into the right flank of C57BL/6J mice. We euthanized tumor-bearing mice on day 15 post-inoculation. We removed and weighed the tumors and cut them into pieces as small as possible. We prepared 1× digestive enzyme solution with 0.2U/ ml Collagenase type IV (Sigma; Cat #C5138) and 50 μg/ml DNase I (Millipore Sigma; SKU #10104159001) in HBSS. We then added the triple enzyme solution to the homogenates and incubated for 1 hour at 37°C to ensure full digestion. We then passed the dissociated cells through a 70 μm cell strainer (BD Falcon; Cat #35-2350) and rinsed them with HBSS three times. The pellets were collected, lysed in 1× red blood cell lysis buffer (Roche; Cat #11814389001) on ice for 5 minutes, and rinsed with 1X wash buffer (2% FBS in 1X PBS). About 1*10^6^ live cells were then blocked with TruStain FcX^TM^ anti-mouse CD16/32 antibody (Clone 93; Biolegend; Cat #101320) on ice for 10 minutes, followed by LIVE/DEAD^TM^ fixable dead cell staining (Thermo Fisher Scientific; Cat #L23105), and cell surface staining using perspective antibodies (see Antibodies and Reagents Section) on ice for 20 minutes. We then fixed the cells with fixation buffer (BioLegend; Cat #420801), permeabilized them with 1× intracellular staining perm wash buffer (Invitrogen; Cat #00-8333-56), and subjected them to intracellular staining using indicated antibodies (See Antibodies and Reagents Section). Samples were analyzed using BD FACS Canto II Flow Cytometer (Flow Cytometry Shared Facility, Duke University School of Medicine).

To evaluate the surface levels of MHC class I in VC and *Vhl*-KO MC38 cells, cells were detached by dissociation buffer (2% EDTA in 1× PBS), followed by blocking with TruStain FcX^TM^ anti-mouse CD16/32 antibody (Clone 93; Biolegend; Cat #101320) on ice for 10 minutes. Cells were then stained with LIVE/DEAD^TM^ fixable dead cell staining (Thermo Fisher Scientific; Cat #L23105) and PE anti-mouse H-2K^b^/H2D^b^ antibody (Clone 28-8-6; Biolegend; Cat #114607) on ice for 20 minutes in the dark, washed twice with 1× FACs buffer (2% FBS in 1×PBS with 2mM EDTA), and fixed with 1% PFA at room temperature for 15 minutes in the dark. Samples were washed twice, resuspended in 1× FACs buffer, and analyzed using BD FACS Canto II Flow Cytometer (Flow Cytometry Shared Facility, Duke University School of Medicine).

### TCR sequencing

We implanted control and *Vhl*-KO MC38 cells into mice as described above. At 13 days post-inoculation, we euthanized the mice and collected tumor tissues, and extracted genomic DNA (gDNA) using QIAGEN^®^ DNeasy Blood & Tissue kit according to manufacturer’s instruction (QIAGEN^®^; Cat # 69506). We sent about 3 μg gDNA (60 ng/μl in AE buffer) to Adaptive Technologies (Seattle, WA) for ImmunoSEQ^®^ mouse TCR-β CDR3 survey sequencing. Upon data acquisition, we analyzed them using ImmunoSEQ^®^ Analyzer 3.0, an Adaptive Biotechnologies online analysis platform.

### Subcellular fractionation of cellular extracts

We followed a previously published procedure for subcellular fractionation of cellular extracts ^46,77^. Briefly, we seeded cells in 100 mm dishes two days before the experiment. On the day of the experiment, we divided 8*10^6^ cells into two equal aliquots. We then suspended one aliquot in 500 μl of 50 mM NaOH, boiled the lysates for 30 minutes and used it to serve as total mtDNA control from whole cell lysate (WCL). Next, we resuspended the other aliquot in 500 μl of solution with 25 μg/ml of digitonin (Millipore Sigma; SKU #D141-100MG) in a buffer with 150mM of NaCl, 50mM HEPES at pH 7.4, and homogenized the cells by vigorous pipetting, and followed by a 10-minute incubation period on ice to allow selective permeabilization of the plasma membrane. We then centrifuged the homogenates at 980 ×g three times for 3 minutes each at 4°C to pellet the intact cells. After the centrifugation, we rinsed the pellet with 1× PBS and used it as the pellet (Pel) fraction for immunoblot. Next, the supernatant was transferred to a new tube and centrifuged at 17,000 ×g for 10 minutes at 4°C to spin down the remaining cellular debris. The supernatant from this centrifugation was transferred to a new tube and used as the cytosolic (Cyto) fraction free of nuclear, mitochondrial, and endoplasmic reticulum contamination. We then boiled the WCL, Pel, and Cyto lysates at 95 °C for 5 minutes and subjected them to western blot analysis to ensure the cytosolic preparation was free of contamination. Finally, we purified and concentrated both WCL_DNA and Cyto_DNA using DNA Clean & Concentrator-5 Kit (ZYMO RESEARCH; Cat #D4004) and used the DNA for quantitative PCR (qPCR) tests.

### Depletion of cellular mtDNA

We depleted mtDNA using ethidium Bromide (EtBr) according to a previously published method ^78,79^. We treated VC and *VHL*-KO cells with 100 ng/mL of EtBr for 21 days with routine medium change every 2-3 days. We then validated the depletion of mtDNA by staining cells with anti-dsDNA and anti-HSP60 antibodies before we harvested the cells for immunoblot and qPCR.

### Total RNA extraction

We plated about 6 *10^5^ VC and *VHL*-KO Caki-1 cells in 60 mm dishes two days before total RNA extraction. We then extracted total RNA using Trizol^®^ Reagent (Ambion by life technologies; Cat #15596018) according to a protocol from StarrLab (https://sites.google.com/a/umn.edu/starrlab/protocols/rna/rna-isolation-using-trizol). Briefly, we first rinsed the cells twice with ice-cold 1× PBS. We then added 1ml Trizol to lyse the cells. Next, we scraped the lysate off from the Petri dish and transferred it to a 1.5 ml Eppendorf tube, incubated the lysate at room temperature for 5 minutes, and added 200 μl chloroform (Sigma; Cat #C2432) with vigorous vortexing for 15 seconds. We then incubated the mixture at room temperature for 10 minutes and centrifuged it at 12,000 ×g for 15 minutes at 4°C. Next, we transferred the transparent top layer to a clean 1.5 ml tube, adding 500 μl isopropanol and incubating at room temperature for 10 minutes. The mixture was then centrifuged at 12,000 ×g for 10 minutes at 4°C to get a white pellet at the bottom. After removing the supernatant, the pellet was washed with 75% ethanol and centrifuged at 7,500 ×g for 5 minutes at 4°C. The pellet was then allowed to air dry and dissolved in 85 μl RNase-free H2O. Subsequently, we added 5 μl of TURBO^TM^ DNase (Invitrogen by Thermo Fisher Scientific; Cat #AM2238) and 10 μl of 10× Turbo DNase buffer to degrade the remaining DNA at 37°C for 30 minutes. We then added 200 μl chloroform to stop the reaction. Afterward, we repeated the RNA extraction procedures as described above. Finally, we dissolved total RNA free of DNA contamination in RNase-free H2O and used it for quantitative PCR and bulk RNA sequencing analysis.

### Quantitative RT-PCR (qRT-PCR) and quantitative PCR

To quantify RNA expression levels, we used the Trizol-extracted total RNA (described above) as the template for cDNA synthesis using random hexamer primers (Invitrogen by Thermo Fisher Scientific; Cat #SO142) and SuperScript II Reverse Transcriptase (Invitrogen Thermo Fisher Scientific; Cat #18064014) following the manufacturer’s instructions. Afterward, we performed qRT-PCR of the cDNA using qPCRBIO SyGreen Blue Mix Hi-ROX (Genesee Scientific; Cat #17-506C) and the Applied Biosystems^®^ ViiA^TM^ 7 Real-Time PCR System with 384-well Block (Thermo Fisher Scientific; Cat #4453536). We used the comparative Ct (ýýCt) method to compare the relative changes in gene expression among different genes.

To quantify cytosolic DNA levels, we obtained cytosolic extracts as described above. We then conducted qPCR analysis using the cytosolic extracts as templates. To quantify cytosolic mitochondria DNA (mtDNA) and nuclear DNA (nucDNA) levels for any individual gene, we set the ratio between the cytosolic DNA level and the whole cell lysate as in the control cells as 1 and used it to obtain the relative levels of cytosolic DNA in other cells.

Table S6 lists the primers used for q-RT-PCR and qPCR analysis of different target genes.

### JC-1 mitochondrial membrane potential assay

We measured mitochondrial membrane potential (MtMP) using the JC-1(tetraethylbenzimidazolylcarbocyanine iodide) mitochondrial membrane potential assay kit (Abcam; Cat #113850) following the manufacturer’s instruction. JC-1 is a lipophilic cationic carbocyanine dye that accumulates in mitochondria. In healthy mitochondria, the mitochondrial membrane is less permeable. The high mitochondrial membrane potential that results from the proton pump causes the aggregation of JC-1, which shows a red to orange color under fluorescence. However, damaged mitochondria are associated with highly permeable and depolarized mitochondrial membrane and reduced MtMP. Under such conditions, JC-1 predominantly forms monomers and produces a green fluorescence. We used FCCP (2-[2-[4-(trifluoromethoxy)phenyl]hydrazinylidene]-propanedinitrile) as a positive control for membrane depolarization as it is an uncoupler of oxidative phosphorylation in the mitochondria ^80^. We assessed the red fluorescence (excitation 535 nm)/emission 590 nm) and green fluorescence (excitation 475 nm/emission 530 nm) using immunofluorescence microscopy and flow cytometry. To quantify green/red fluorescence ratios in flow cytometry, the following formula was used: Ratio = [Red^+^Green^+^ (Q2, quadrant 2) + Red^-^ Green^+^ (Q3)] / Red^+^Green^-^ (Q1).

### Immunofluorescence microscopy

We seeded the cells in 35 mm glass-bottomed poly-D-lysine-coated dishes (MatTek^TM^ Life Sciences; Cat #P35G-1.5-10-C) two days before experiments. We used 4% Paraformaldehyde (PFA) to fix cells at room temperature for 15 minutes and permeabilized the cells using 0.5% Triton X-100 in PBS at room temperature for 10 minutes. We then washed the cells three times with PBS and blocked them with 5% bovine serum albumin (BSA; Sigma; Cat #A3983) at room temperature for 1 hour. Next, we added primary antibodies and incubated the cells at 4°C overnight, followed by adding fluorophore-conjugated secondary antibodies after washing with PBS three times. Next, we incubated the cells at room temperature for 1 hour in the dark and washed them three times. Finally, we added VECTASHIELD^®^ Antifade Mounting Medium with DAPI (VECTOR LABORATORIES; Cat #H-1200-10) to the glass bottom of the dish before analysis. We took fluorescence images using the Leica TCS SP5 laser scanning confocal microscope in the Light Microscopy Core Facility of Duke University School of Medicine.

### Bulk RNA sequencing

To perform genome-wide transcriptome analysis of VC and *VHL*-KO Caki-1 cells, we prepared total RNAs from the cells using Trizol^®^ Reagent as described above. We then submitted our RNA samples to the Duke Center for Genomic and Computational Biology for sequencing, which QC’ed the samples and prepared cDNA libraries for analysis using Illumina NovaSeq 6000. We processed the RNA-seq data using the TrimGalore toolkit, which employs Cutadapt to trim low-quality bases and Illumina sequencing adapters from the 3’ end of the reads ^81,82^. Only reads that were 20nt or longer after trimming were kept for further analysis. Reads were mapped to the GRCh38.p13 of the human genome and transcriptome using the STAR RNA-seq alignment tool ^83,84^. Reads were kept for subsequent analysis if they mapped to a single genomic location using the SAMtools ^85^. Gene counts were compiled using the HTSeq tool^86^. Only genes that had at least 10 reads in any given library were used in subsequent analysis. Normalization and differential expression were carried out using the DESeq2 Bioconductor package with the R statistical programming environment. Software for gene set enrichment analysis (GSEA; version 4.1.0) was used to identify differentially regulated pathways. The source RNA-seq data are deposited in the NCBI’s Gene Expression Omnibus (GEO) database.

### Analysis of published RNAseq data

To assess enriched pathways from VHL overexpression in 786-O cells, a previously published dataset (GSE108229) was used to perform gene ontology (GO) analysis ^87^. Top-ranked pathways were plotted with ‘ggplot2’ R package. Scripts are available upon request.

### Human data analysis

All human data we obtained are publicly available. To study whether *VHL* mutations enhance ICB therapy in ccRCC patients, we integrated three cohorts of ICB-treated ccRCC patients^26,28,61^ and compared the clinical outcomes (overall survival, OS) of patients with wild-type *VHL* or *VHL* mutations. In the CheckMate 009/010/025 cohorts, only patients with available RNA sequencing data were included. The group identified as WT *VHL* were those patients without any *VHL* mutations. Those with *VHL* mutations were identified by genetic alternations. The OS of two groups were compared through Kaplan-Meier survival analysis (GraphPad Prism version 8.2.0). We also used these data to analyze other frequently mutated genes in ccRCC and their impact on ICB therapy. The OS of patients with mutations of *PBRM1, SETD2, BAP1*, or *KDM5C* were compared to patients with respective wild-type genes.

To compare the clinical outcomes (overall survival, OS) between RCC patients with WT *VHL* and those with *VHL* mutations, we accessed ten PanCancer studies from the open-access online tool cBioPortal (http://www.cbioportal.org) ^88,89^. These studies include kidney renal clear cell carcinoma (KIRC), kidney renal papillary cell carcinoma (KIRP), and kidney chromophobe carcinoma (KICH) patients from the Cancer Genome Atlas (TCGA) Pan-Cancer studies ^90–92^, and nine PanCancer Studies available to the date of analysis (Nov 28, 2023) ^93–101^.

To identify patients with *VHL* mutations across multiple cancers, we listed *VHL* mutation ratios (Ratio = patient cases with *VHL* mutations / total cases). Data were collected from three cohorts, including MSK-IMPACT (Table S1) ^102^, China-PanCancer (Table S2), and TCGA PanCancer (Table S3) ^90–92^. We also combined nine immunogenomic studies to further evaluate the association of *VHL* mutations with ICB therapy across different cancers, including glioblastoma, melanoma, lung cancers, and bladder cancers ^26,28,61–67^.

### Statistical analysis

Statistical analysis was conducted using GraphPad Prism 8.2.0 software. Two-sided Student’s *t* test was used for comparing two experimental groups. One-way ANOVA was applied to compare gene expression levels among multiple groups. Two-way ANOVA was applied to compare *in vivo* tumor growth rates within two or more experimental groups. mtDNAs and nucDNAs levels in VC, *VHL*-KO, *VHL/HIF1α* DKO, *VHL/HIF2α* DKO, and *VHL/HIF1α/HIF2α* TKO Caki-1 cells were also analyzed using two-way ANOVA. Log-rank (Mantel-Cox) test was used for mouse and human patient survival analysis. \**P* < 0.05 was considered statistically significant.

## Data Availability

Raw RNAseq and metadata are deposited in the Gene Expression Omnibus database with the accession number GSE196509. Source data for other figures will also be provided.

## Author Contributions

M.J. and C.-Y.L. designed the study. M.J. carried out CRISPR-Cas9-mediated gene knockouts in tumor cells. M.J. D.P. and M.H. performed western blot analysis. M.J., J.K., M.H., and D.P. carried Q-RT-PCR analysis. M.J. carried out tumor growth experiments. M.J. and X.B. characterized tumor cells in vitro and in vivo and intratumoral lymphocytes in vivo using flow cytometry. M.J. and X.B. carried out RNAseq analysis. X.L. and F.L. advised on CRISPR knockouts. F.L. provided material support. M.J. and C.-Y.L. wrote the manuscript with help from all co-authors. C.-Y.L. provided funding and study supervision.

## Acknowledgments

We thank J.M. Cook and colleagues at the Flow Cytometry Facility of Duke University School of Medicine for their expert assistance. We further thank the Duke University Light Microscopy Core Facility for professional help with confocal microscopy. Our study is supported in part by US National Institutes of Health (NIH) grants R01CA208852, R01CA216876, R01CA251439, and P30CA014236.

---

**Figure S1.**
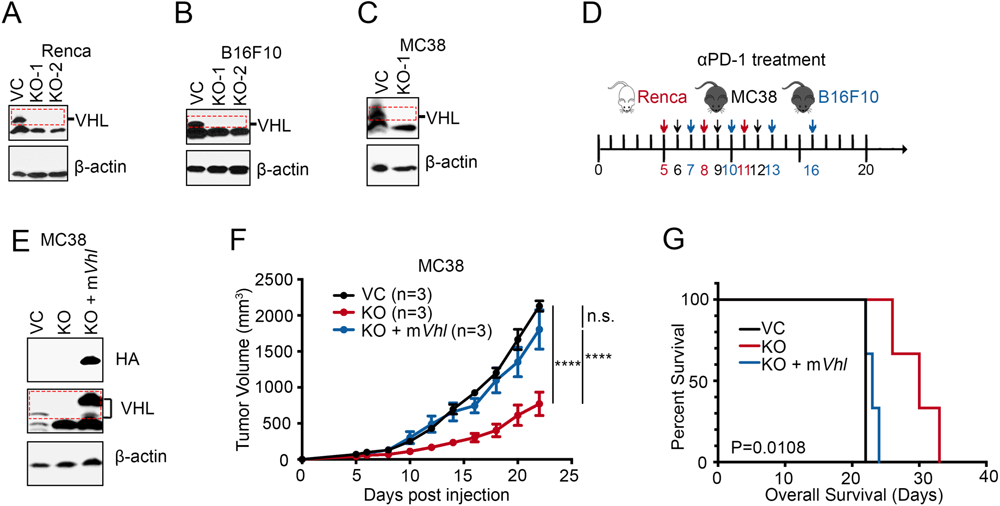
Additional data on the role of *Vhl* gene loss in tumor growth. **(A-C)** Immunoblot showing genomic knockout of *Vhl* in Renca (A), B16F10 (B), and MC38 (C) cells. **(D)** Scheduling of for αPD-1 antibody treatments in mice bearing Renca, B16F10, and MC38 tumors. **(E)** Immunoblot verification of the re-expression of wild-type mouse *Vhl (mVhl)* in *Vhl*-KO MC38 cells. **(F-G)** Tumor growth (F) and Kaplan-Meier survival curves (G) of C57BL/6J mice subcutaneously implanted with 5x105 VC, *Vhl*-KO, and *Vhl*-KO MC38 cells re-expressing m*Vhl* (n=3). Error bars represent mean ± SEM. *P<0.05; **P<0.01; ***P<0.001; ****P<0.0001; n.s. not significant; two-way ANOVA.

**Figure S2.**
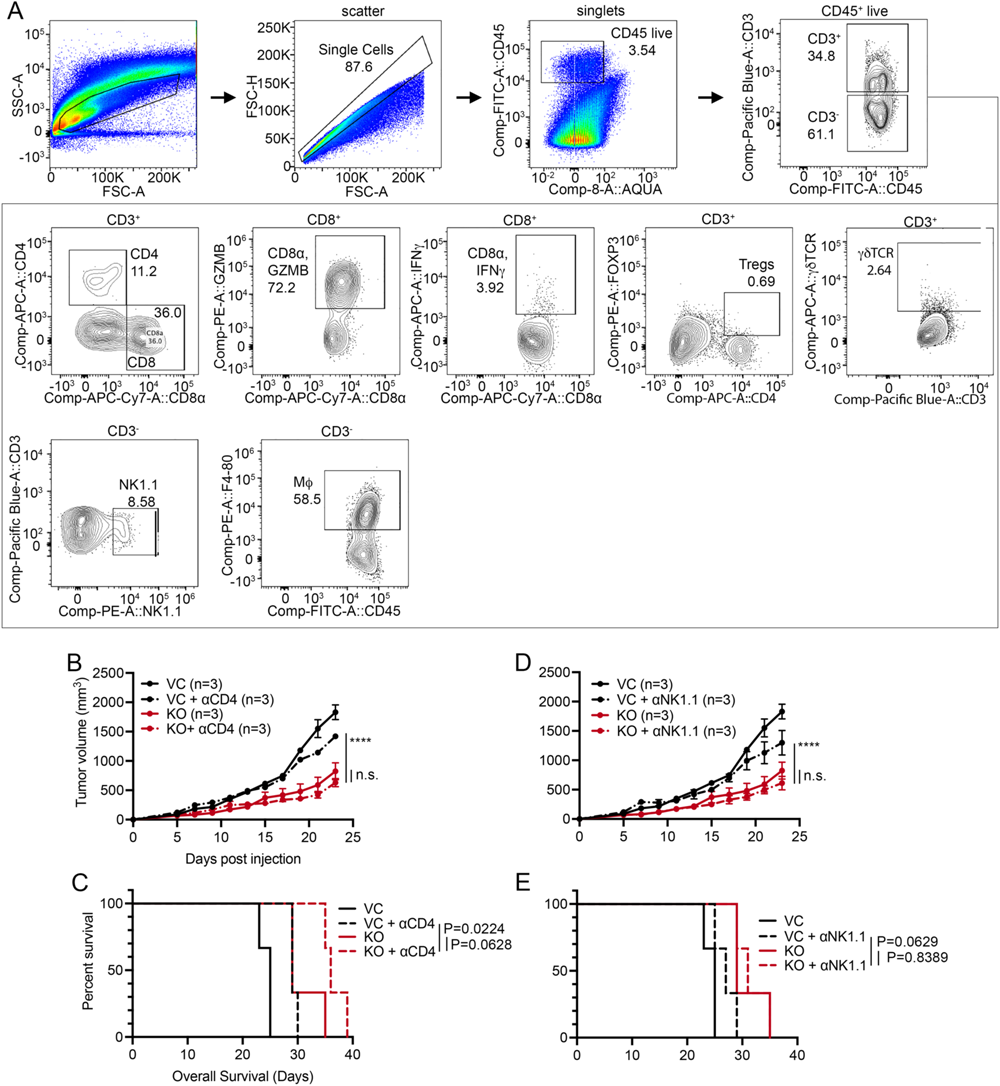
Additional data on the involvement of immune effector cells in *Vhl* loss mediated tumor growth delay. **(A)** Gating strategy of TIL analysis by flow cytometry. **(B-C)** Tumor growth (B) and Kaplan-Meier survival curves (C) of C57BL/6J mice bearing VC and *Vhl*-KO MC38 tumors in mice injected with isotype or αCD4 antibodies (n=3). **(D-E)** Tumor growth (D) and Kaplan-Meier survival curves (E) of C57BL/6J mice bearing VC and *Vhl*-KO MC38 tumor injected with isotype or αNK1.1 antibodies (n=3). In B and D, error bars represent mean ± SEM. *P<0.05; **P<0.01; ***P<0.001; ****P<0.0001; n.s. not significant; two-way ANOVA. In C and E, p values calculated by using the log-rank test.

**Figure S3.**
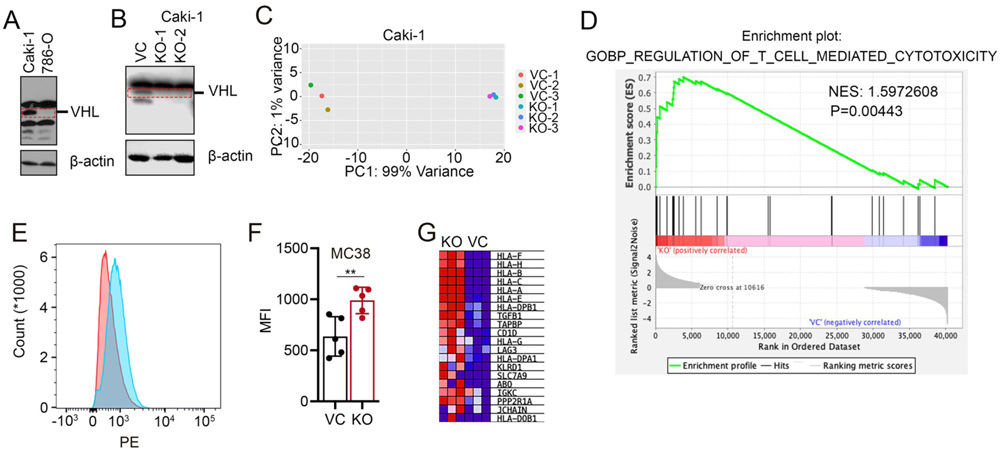
Supporting data of control and *VHL*-KO Caki-1 cells and comparison of surface MHC-I expression in Caki-1 cells. **(A)** Immunoblots of *VHL* expression in Caki-1 and 786-O cells. **(B)** Immunoblot analysis verifying the knockout of *VHL* in Caki-1 cells. **(C)** PCA analysis of RNA sequencing data generated from control and *VHL*-KO Caki-1 cells. **(D)** GSEA analysis indicating the enrichment of regulation of T cell mediated cytotoxicity in *VHL*-KO Caki-1 cells. (**E-F)** Flow cytometry analysis of the levels of H2Kb/H2Db in VC and *Vhl*-KO MC38 cells. Data in B from five independent experiments. **(G)** Heatmap of top-20 differently expressed genes involved in antigen binding in *VHL*-KO and control Caki-1 cells. Error bars represent mean ± SD. *P<0.05; **P<0.01; ***P<0.001; ****P<0.0001; n.s. not significant.

**Figure S4.**
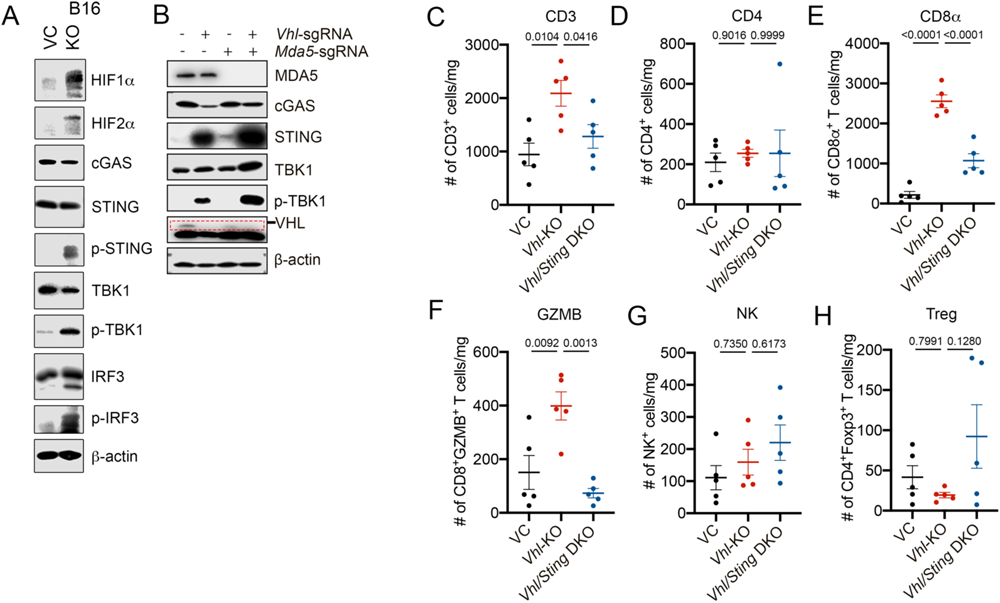
The importance of cGAS-STING mediated DNA sensing but not MDA5-mediated RNA sensing pathway in *Vhl*-KO induced anti-tumor immune responses. **(A)** Upregulation of cGAS-STING signaling and its downstream effectors in B16F10 cells with *Vhl* gene loss. **(B)** Immunoblot analysis of cGAS-STING and type I interferon signaling pathway in VC, *Vhl*-KO, *Mda5*-KO, and *Vhl*/*Mda5* DKO MC38 cells. **(C-H)** Tumor infiltrated lymphocytes from VC, *Vhl*-KO, and *Vhl/Sting* DKO MC38 tumors by flow cytometry (n=5 per group). Error bars represents mean ± SEM.

**Figure S5.**
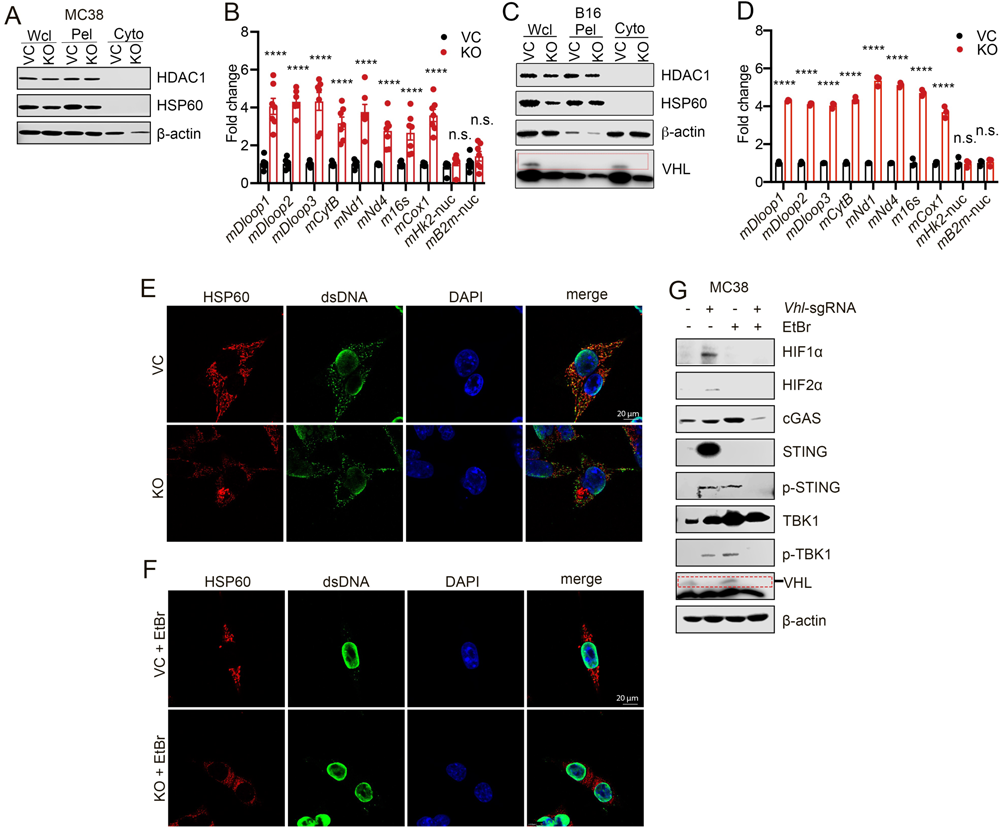
Additional data on *Vhl* loss induced mtDNA leakage. (A-D) Comparison of cytosolic mtDNA levels in VC and *Vhl*-KO cells. (A, C) Immunoblot analysis validating the purity of cytosolic extracts from VC and *Vhl*-KO MC38 (A) and B16F10 (C) cells. (B, D) Quantitative PCR analysis of cytosolic mtDNAs in VC and *Vhl*-KO MC38 (B) and B16F10 (D) cells. Results from three independent experiments. **(E)** Immunofluorescent microscopy of VC and *VHL*-KO MC38 cells stained with anti-HSP60 and anti-dsDNA antibodies. Scale bar, 20μM. **(F)** Immunofluorescent microscopy verifying mtDNA depletion in VC and *Vhl*-KO MC38 cells treated with 100 ng/ml EtBr. Scale bar, 20μM. **(G)** Immunoblot analysis of cGAS-SITNG signaling in VC and *Vhl*-KO MC38 cells treated with vehicles or 100 ng/ml EtBr. Error bars represent mean ± SEM. *P<0.05; **P<0.01; ***P<0.001; ****P<0.0001; n.s. not significant.

**Figure S6.**
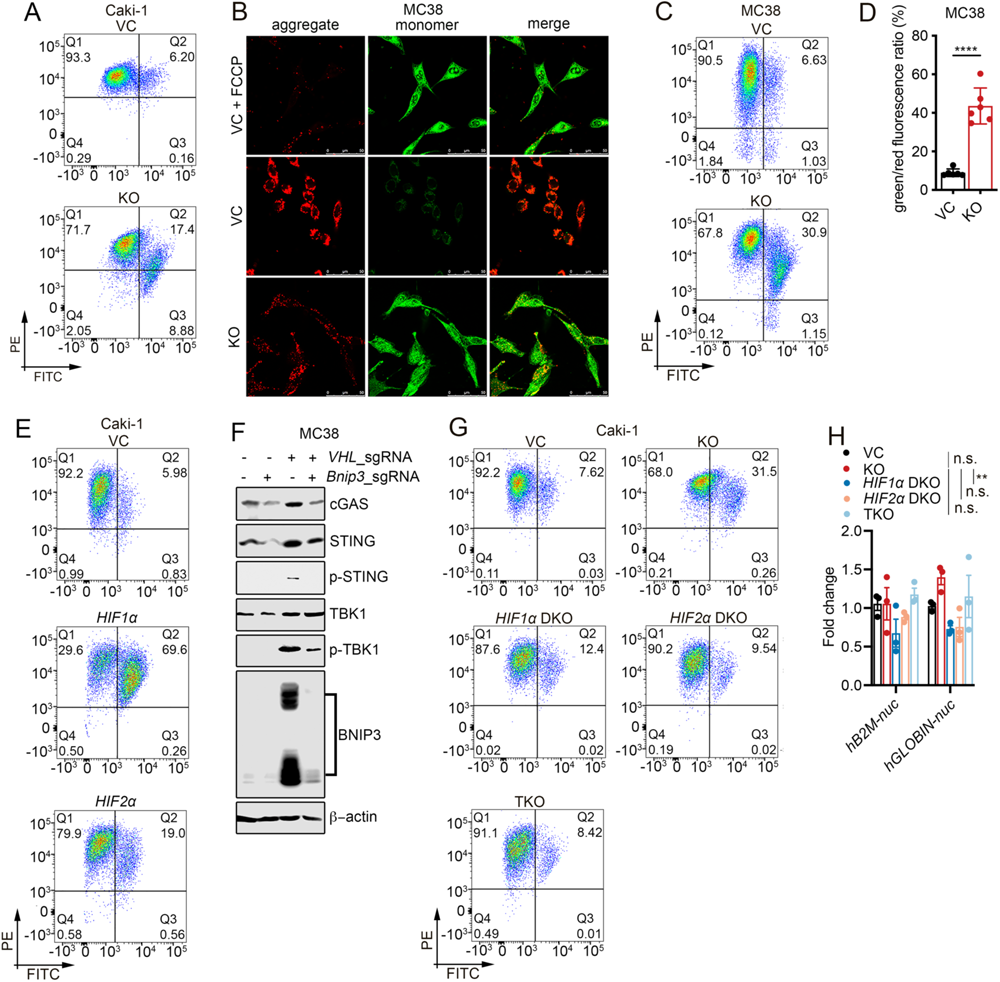
Additional data demonstrating that HIF1α and HIF2α are functionally required for the *VHL* deficiency-induced mitochondrial membrane permeabilization. **(A)** Flow cytometry of JC-1 staining of VC and *VHL*-KO Caki-1 cells. **(B-D)** JC1-enabled mitochondrial membrane potential (MtMP) assay for VC and *Vhl*-KO MC38 cells as detected by confocal microscopy (B) and flow cytometry (C-D). Scale bar, 50μM. Error bars represent mean ± SD. **(E)** Flow cytometry analysis JC-1 staining of VC, degradation-resistant mut*HIF1α* (h*HIF1α*-p402A/p564A), and mut*HIF2α* (h*HIF2α*-p405A/p531A) expressing Caki-1 cells. **(F)** Immunoblot analysis of cGAS-STING signaling proteins in VC, *Bnip3*-KO, *Vhl*-KO, and *Vhl*/*Bnip3* DKO MC38 cells. **(G)** Flow cytometry analysis of JC-1 staining in VC, *VHL*-KO, *VHL*/*HIF1α* DKO, *VHL*/*HIF2α* DKO, and *VHL*/*HIF1α*/*HIF2α* TKO Caki-1 cells. **(H)** Quantitative PCR analysis of cytosolic nucDNAs in VC, *VHL*-KO, *VHL*/*HIF1α* DKO, *VHL*/*HIF2α* DKO, and *VHL*/*HIF1α*/*HIF2α* TKO Caki-1 cells. Data from three independent experiments. Error bars represent mean ± SEM. Two-way ANOVA. *P<0.05; **P<0.01; ***P<0.001; ****P<0.0001; n.s. not significant.

**Figure S7.**
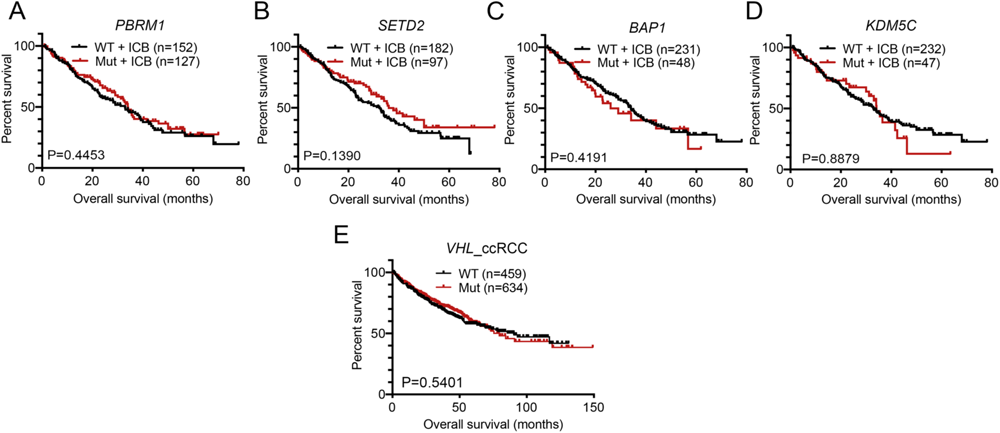
Additional survival analysis of human RCC patients with different gene mutations. **(A-D)** Kaplan-Meier survival analysis of ICB-treated ccRCC patients with or without *PBRM1* (A), *SETD2* (B), *BAP1* (C), and/or *KDM5C* (D) gene mutations in combination with ICB therapy. **(E)** Kaplan-Meier survival analysis of non-ICB treated ccRCC patients who are WT or with *VHL* mutations from PanCancer cohorts on cBioPortal.

**Table S1:**
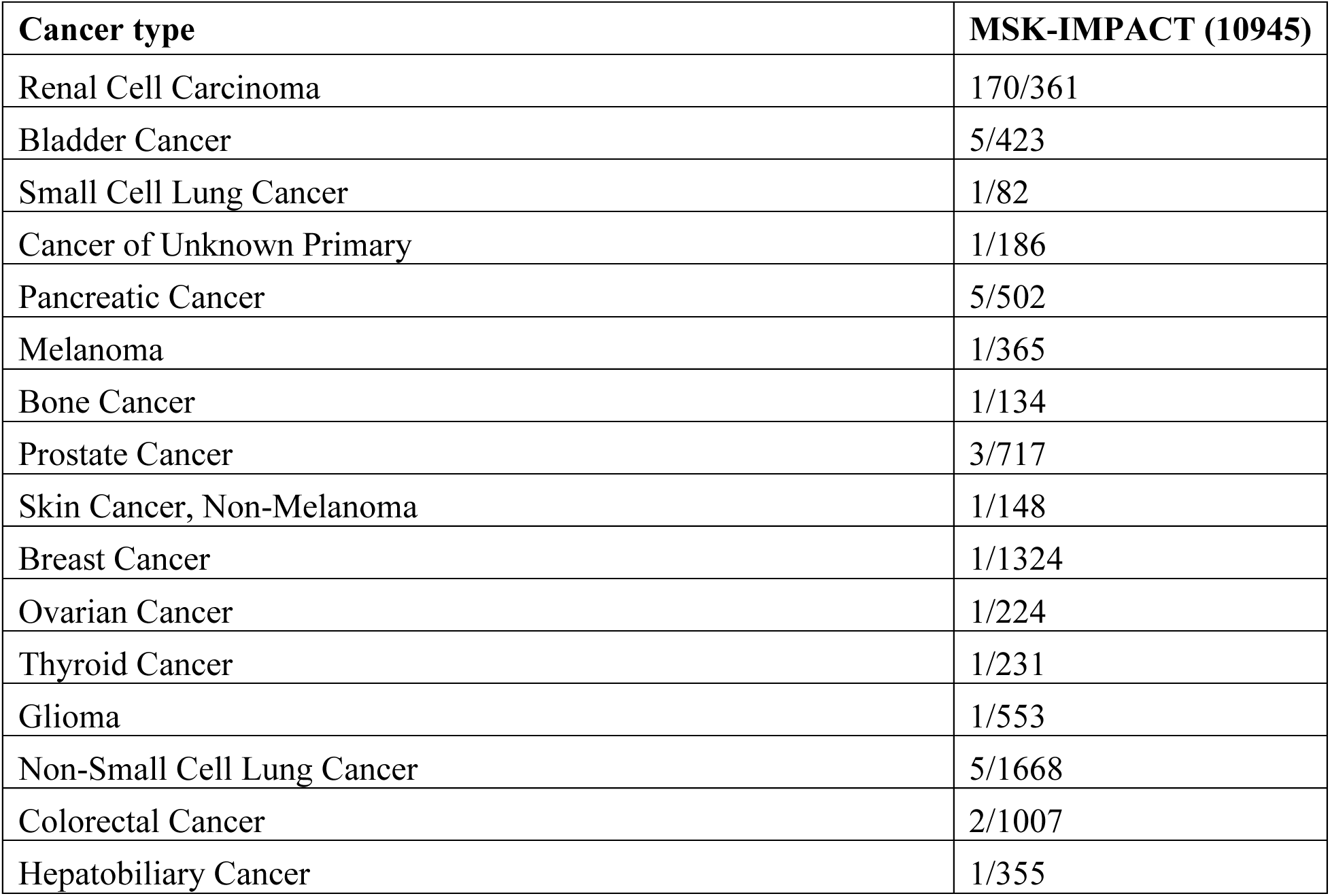
Patient cases with VHL mutations in MSK-IMPACT cohort.

| **Cancer type** | **MSK-IMPACT (10945)** |
| --- | --- |
| Renal Cell Carcinoma | 170/361 |
| Bladder Cancer | 5/423 |
| Small Cell Lung Cancer | 1/82 |
| Cancer of Unknown Primary | 1/186 |
| Pancreatic Cancer | 5/502 |
| Melanoma | 1/365 |
| Bone Cancer | 1/134 |
| Prostate Cancer | 3/717 |
| Skin Cancer, Non-Melanoma | 1/148 |
| Breast Cancer | 1/1324 |
| Ovarian Cancer | 1/224 |
| Thyroid Cancer | 1/231 |
| Glioma | 1/553 |
| Non-Small Cell Lung Cancer | 5/1668 |
| Colorectal Cancer | 2/1007 |
| Hepatobiliary Cancer | 1/355 |

**Table S2:** Patient cases with VHL mutations in China PanCancer cohort.

| **Cancer type** | **China PanCancer (10194)** |
| --- | --- |
| Renal Cell Carcinoma | 177/308 |
| Small Cell Lung Cancer | 1/220 |
| Breast Cancer | 1/323 |
| Ovarian Cancer | 1/261 |
| Non-Small Cell Lung Cancer | 7/2039 |
| Colorectal Cancer | 9/1225 |
| Hepatobiliary Cancer | 6/1133 |
| Gastric Cancer | 2/866 |
| Esophageal Carcinoma | 1/610 |
| Head and Neck carcinoma | 1/175 |
| Intrahepatic Cholangiocarcinoma | 1/555 |
| Gastrointestinal Neuroendocrine Tumor | 1/74 |

**Table S3:** Patient cases with VHL mutations in PanCancer cohort.

| **Cancer type** | **PanCancer (10953)** |
| --- | --- |
| Kidney Renal Clear Cell Carcinoma | 159/511 |
| Bladder Urothelial Carcinoma | 2/411 |
| Prostate Adenocarcinoma | 3/494 |
| Thymoma | 2/123 |
| Pheochromocytoma and Paraganglioma | 3/178 |
| Stomach Adenocarcinoma | 3/440 |
| Skin Cutaneous Melanoma | 5/444 |
| Kidney Chromophobe | 1/65 |
| Uterine Corpus Endometrial Carcinoma | 5/529 |
| Kidney Renal Papillary Cell Carcinoma | 3/283 |
| Colorectal Adenocarcinoma | 4/594 |
| Lung Squamous Cell Carcinoma | 1/487 |
| Lung Adenocarcinoma | 1/566 |

**Table S4:**
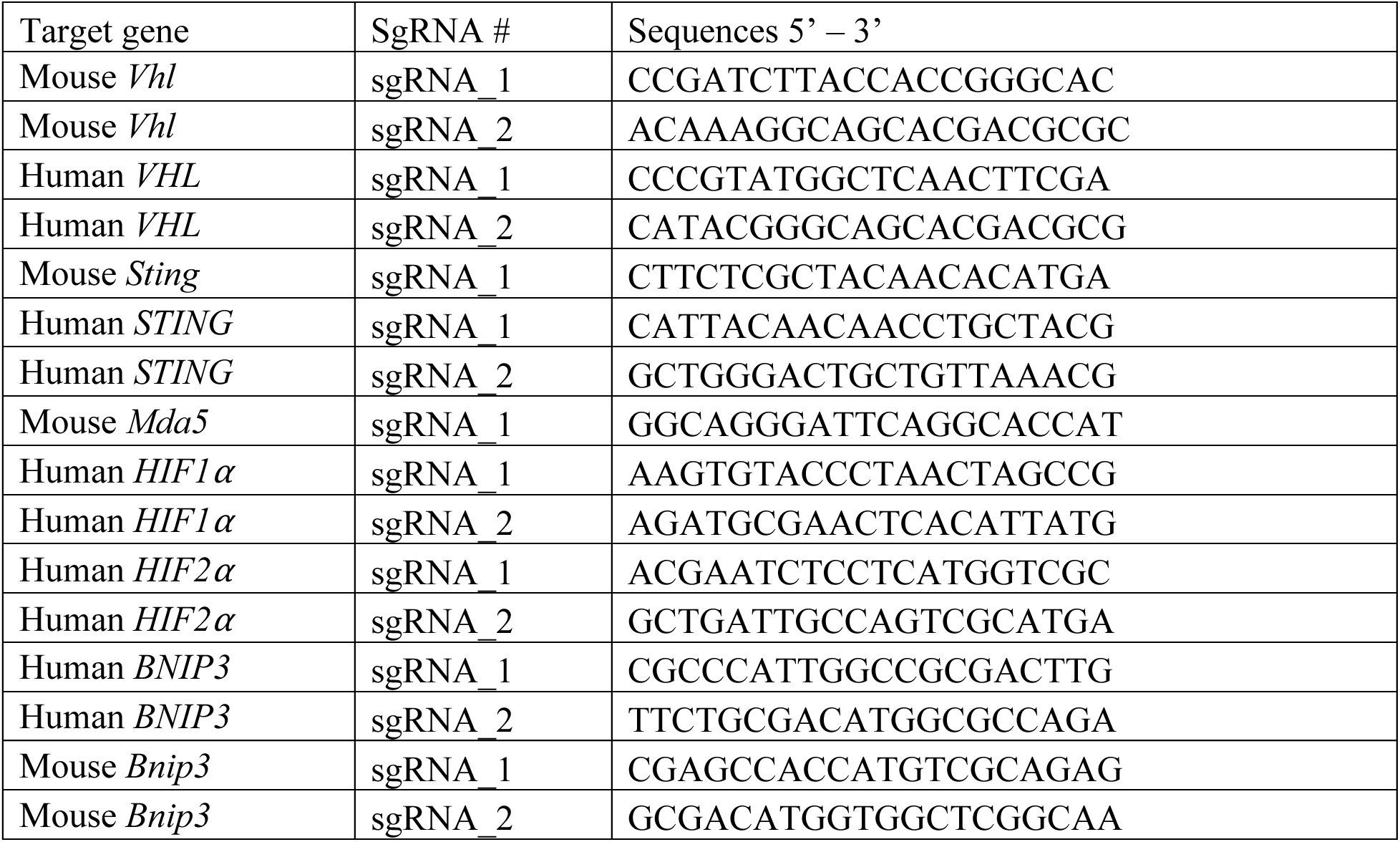
sgRNAs for CRISPR/Cas9 mediated gene knockout.

**Table S5:**
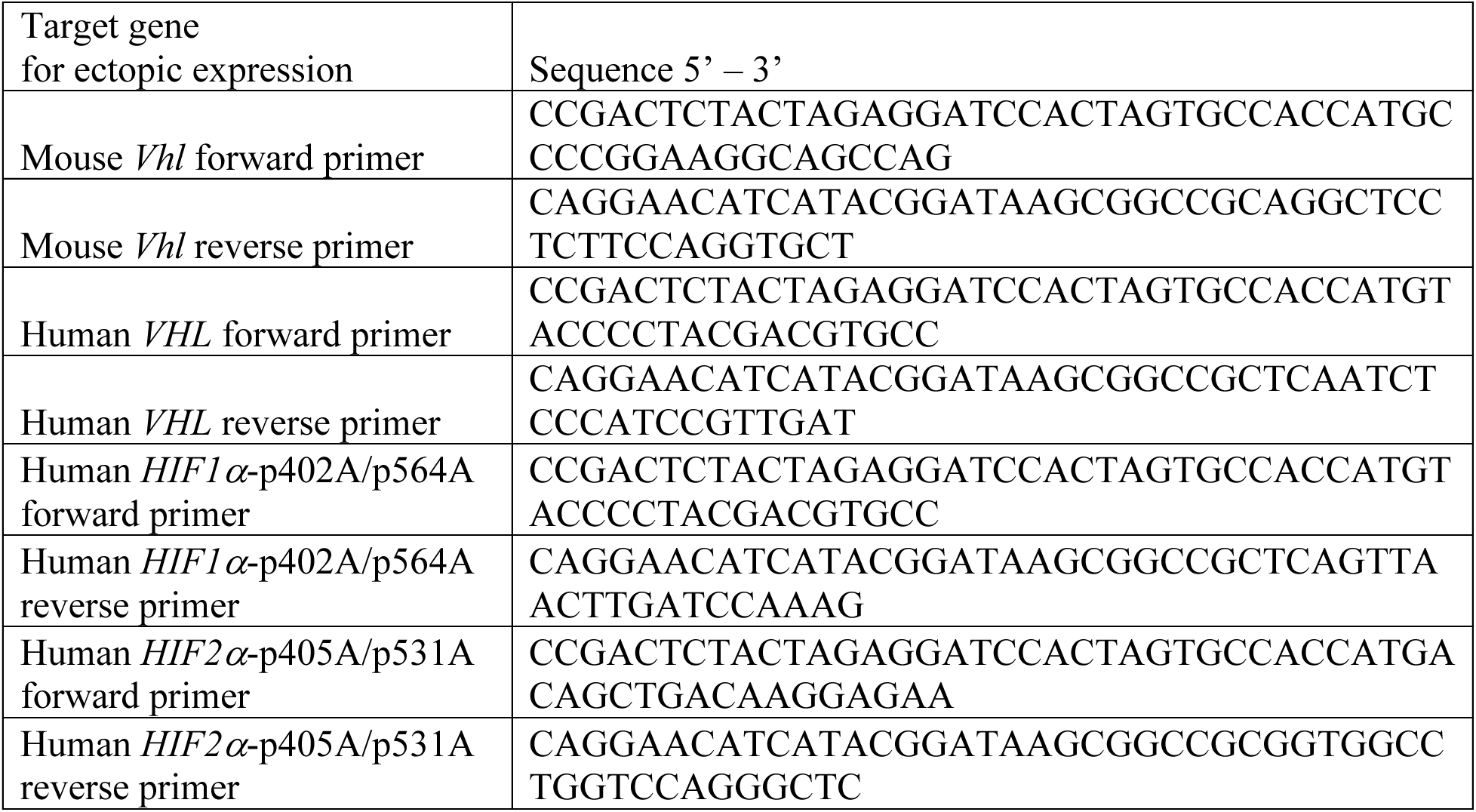
Primers for ectopic gene expression.

**Table S6:**
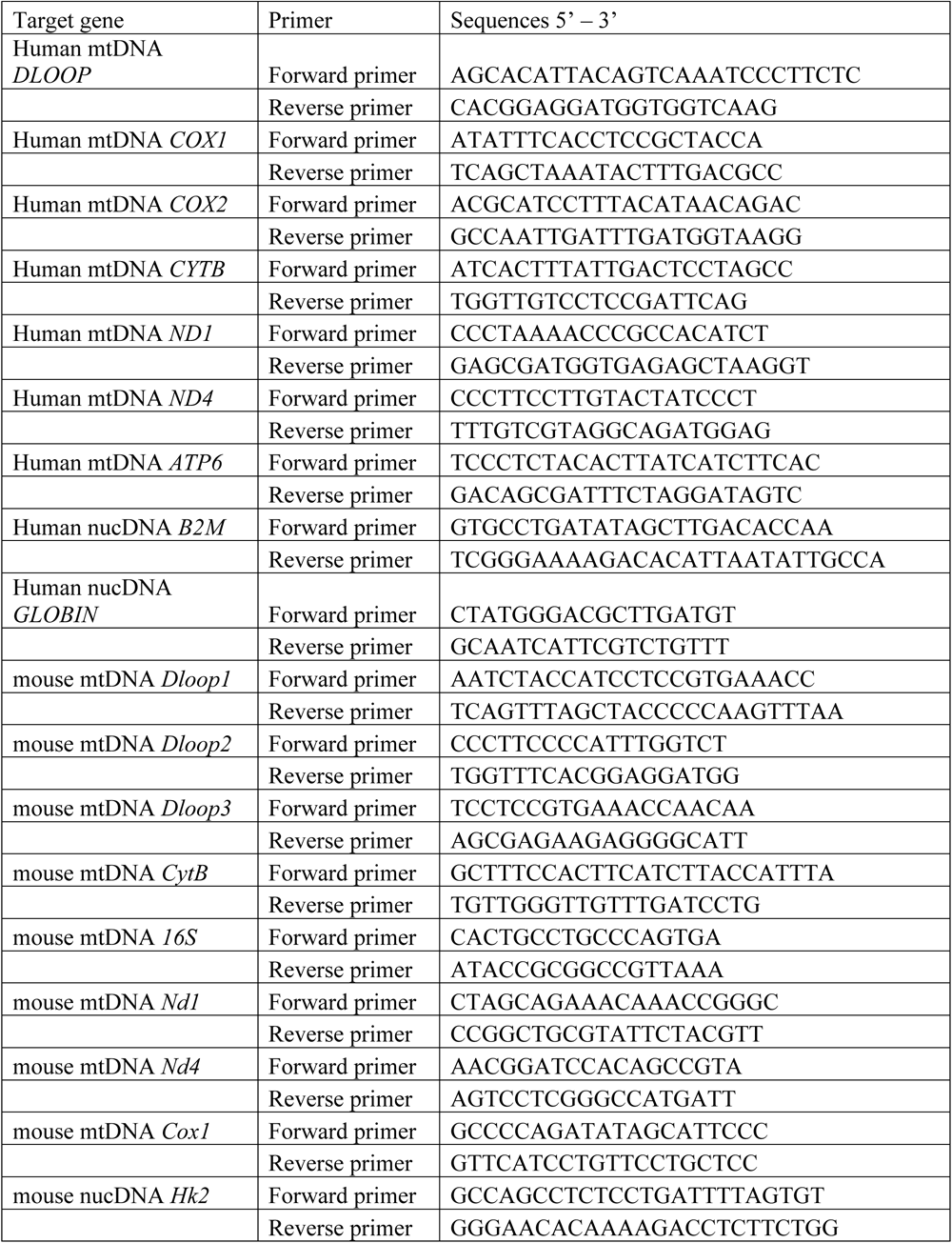
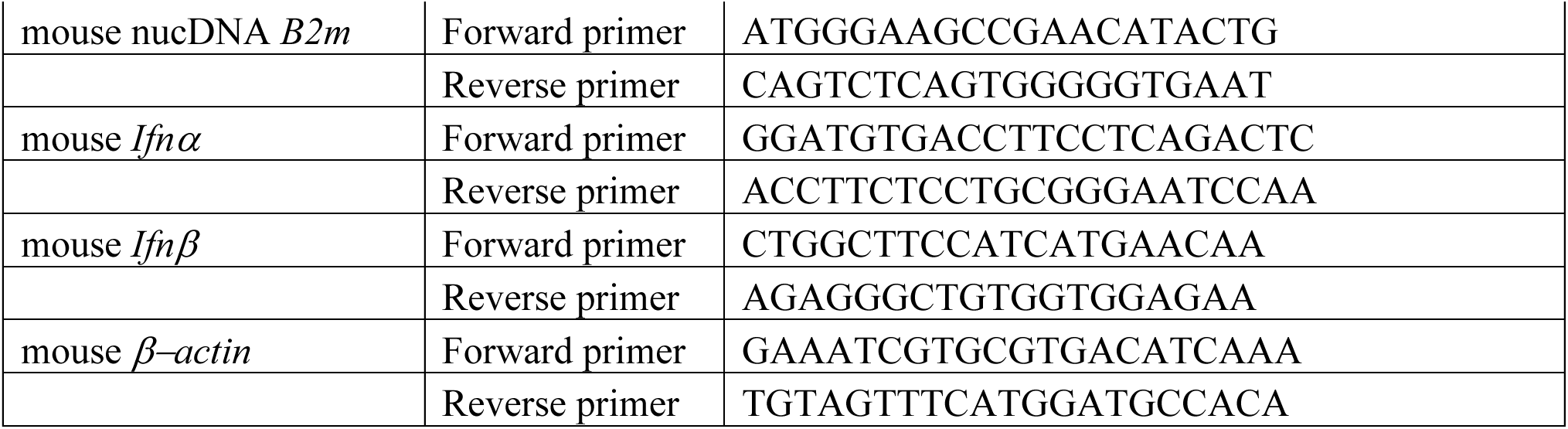
Primers for quantitative RT-PCR and quantitative PCR.

